# Negative frequency-dependent selection contributes to modular structure of effector repertoires in *Pseudomonas syringae*

**DOI:** 10.1101/2025.01.16.632277

**Authors:** Shuanger Li, Eric Laderman, Hanna Märkle, Yunze Yang, Joy Bergelson, Mercedes Pascual

## Abstract

The evolutionary fate of multi-strain pathogens is shaped by host-pathogen ecological interactions. In bacterial pathogens of plants, enhanced strain characterization and advances in our understanding of molecular mechanisms underlying defense pathways open the door for revisiting the role of negative frequency-dependent selection (NFDS) in strain structure, including its interplay with genetic exchange. NFDS arising from specific defense is one potential mechanism for generating, maintaining, and structuring pathogen diversity. In plants, specific protection against microbial pathogens involves Resistance proteins (R-proteins) that recognize virulence factors (effectors) secreted by pathogens, typically to subvert the initial line of host defense. Here we formulate a stochastic computational co-evolution model that explicitly incorporates variable length R-gene and effector repertoires, and migration from their regional pools. We use this model to understand potential mechanisms shaping effector repertoire structure and associated strain coexistence in the generalist plant pathogen *P. syringae*. The demonstration of a modular structure in our numerical simulations motivates the analysis of genome sequences from 76 strains collected in the Midwestern US and 1104 strains from global sources. We find that effector repertories both locally and globally exhibit a modular structure, with higher similarity within than between clusters. The observed modules are consistent with the core genome phylogeny and are unexplained by plant host species, location of isolation, and genetic linkage between effectors. An extension of the model is needed to take into account the evidence for genetic exchange and the phylogenetic congruence of effector modules. We initialize the system with a phylogenetically congruent modular structure and include recombination rates decreasing as a function of phylogenetic distance. We show that NFDS can counter-balance the effects of mixing due to recombination and in so doing, contributes to the maintenance of strain structure. These findings indicate that the observed similarity clusters may constitute, in part, emergent niches arising from eco-evolutionary dynamics that contribute to strain coexistence.

## 1 Introduction

With their extensive intra-specific genetic variation and fast generation times, microbial populations present both an opportunity and a challenge to address the interplay of ecology and evolution in shaping strain coexistence and the maintenance of strain diversity [95, 94, 93, 98, 21, 68, 88]. Of particular relevance is the role played by ecological interactions mediated through specific traits that are advantageous when rare and disadvantageous when common, leading to negative frequency-dependent selection (NFDS) [6, 25, 61, 43]. The intra-specific diversity of microbial populations can span large phylogenetic distances and different degrees of genetic exchange [39, 40]. The extent to which NFDS structures the diversity of strains of human pathogens, especially in the face of mixing due to recombination and horizontal gene transfer (HGT), is an active area of research related to acquired immunity and cross-immunity [45, 7, 43]. Understanding the maintenance of the resulting large diversity is important due to its potential consequences for pathogen resilience [44, 102].

Plants do not have specific adaptive immunity; instead, they employ nucleotide-binding leucine-rich repeat (NLR) proteins, encoded by Resistance genes, or R-genes, that recognize virulence factors of microbes known as effectors, either directly or indirectly, and initiate a defense response upon successful recognition [30, 8, 48, 24, 23]. Pathogens secrete effectors to subvert plant general immunity that targets relatively conserved microbial patterns [23]. Effectors that interfere with various host innate immunity pathways have been identified in a wide range of pathogens, including bacteria, oomycetes, and fungi [13, 54]. Pathogens suffer from NLR recognition, which they may escape via mutation of effectors or the absence of a cognizant NLR-gene in the plant [62, 15]. The evasion of NLR recognition whilst maintaining a virulence function confers a fitness advantage to pathogen strains, whereas successful recognition of effectors benefits the plant. Together, these interactions can drive frequency fluctuations of effectors and R-genes [91, 58, 48]. Thus, plant pathogens represent another large class of microbes in which to investigate the role of NFDS in shaping strain structure and diversity, but in a context of host recognition and defense other than that of acquired immunity. Here, we consider strain diversity from the perspective of effector repertoires.

*Pseudomonas syringae* is one of the most widespread and well-studied plant bacterial pathogens, for which thousands of effector alleles have been identified from hundreds of strains [32, 59]. Most of these effectors are found in the accessory genome and are patchily distributed across the *P. syringae* phylogeny [9, 33, 32]. Single strains carry from a few to about fifty effectors, which creates a large space of possible repertoires with diverse virulence functions to evade plant immunity [33, 31]. *P. syringae* is a generalist pathogen, and currently has been classified into over sixty pathovars [9]. Notably, the pathovar classification is not necessarily congruent with phylogenetic relatedness [86, 84]; one possible contributing mechanism is that genetic exchange alters the distribution of effectors, and therefore pathogenicity, across *P. syringae* phylogeny [86]. Genetic exchange can rearrange accessory genes of diverse functions and backgrounds, and facilitate the spread of beneficial genes [55, 57].

The “ecotype” model, a conceptual model to explain persistent bacterial diversity, proposes that diversification results in lineages occupying distinct ecological niches. Strains sharing the same resource are derived from the same lineage and can be grouped into ecotypes [26]. This model assumes that homologous recombination promotes sweeps within ecotypes but becomes less frequent as sequence divergence between strains increases. This restricted gene flow serves to fortify the lineage structure while retaining cohesiveness [26, 27, 56, 40, 60]. Importantly, the concept of niches in the ecotype model relies on established differences in strains’ life-history (ex. environmental versus pathogenic) or the host (ex. pathovars).

This conceptual view differs from one in which niches emerge intrinsically from the population dynamics of transmission and defense, growth and recognition, as the result of variation in fitness purely as a function of frequency. NFDS can lead dynamically to the emergence of subgroups of strains whose effector phenotypes are more similar within than between clusters. These subgroups would respectively rely for their growth on different subgroups of plants defined in terms of R-gene similarity, exploiting in this way hosts less able to recognize their constituent strains. Thus, NFDS can generate population structure within both interacting partners, with emergent niches representing different subgroups in the host population, differentiated by their defense against pathogens. A large body of host-pathogen models has sought to understand the contribution of negative frequency-dependent selection to the maintenance of allelic diversity in the context of “gene-for-gene” and “matching allele” frameworks[1, 42, 37, 70, 75], largely formulated before the understanding of underlying molecular mechanisms. For analytical tractability, their formulation has often focused on a small and fixed set of interacting loci, and thus on a fixed length of host resistance genes and pathogen effector repertoires. The increasing availability of host and pathogen genomic data has revealed variable numbers of resistance and effector genes within genomes, opening the door for further interrogating the specific structure of these repertoires and the role of negative-frequency dependent selection in shaping such structure in nature[15, 41].

Here, we develop a stochastic computational model for the temporal dynamics of a plant-pathogen system, to address the associated diversity and structure of plant R-gene and pathogen effector repertoires. Our model is rooted in classic multi-locus gene-for-gene models [85, 5], but does not focus on a fixed number of static loci with virulent/avirulent and susceptible/resistance alleles. Rather, we allow for variably sized R-gene and effector repertoires for each individual and incorporate sampling from a given regional pool. This formulation allows for a simple representation of presence/absence patterns of effectors found in empirical observations [49, 74, 31], while also providing the possibility of incorporating a mechanism for genetic exchange. With numerical simulations, we investigate patterns of strain coexistence and associated population structure in terms of effector repertoire similarity. For comparison, we analyze local strain structure empirically from genomic data for *P. syringae* sampled in a region of the Midwestern US, and further take advantage of existing global sequence datasets. We find evidence of a modular structure in these data, consistent with the role of selection in our initial model. Two additional observations are not accounted for by the model: the modules are found congruent with the phylogeny of the pathogen, and the subfamily distribution of effectors across genomes is consistent with genetic exchange (see also [33]). We therefore modify the model to explore the role of NFDS in maintaining a modular strain structure despite the homogenizing roles of genetic exchange. We discuss this structure in the context of host niches and other ecological systems in which frequency or density-dependent interactions play a role in diversity and coexistence.

## 2 Results

### 2.1 Computational model of plant-microbial eco-evolutionary dynamics generates a modular strain structure

Motivated by the *A. thaliana* and *P. syringae* system, we developed a discrete-time stochastic computational model where plant and pathogen genotypes are characterized by their R-gene and effector repertoires, respectively (for details, see Methods 6.4). Time is discrete to represent generations of an annual plant. The model explicitly tracks the infection status and the genotype of individual plants to take into account demographic stochasticity and to explicitly follow the large number of genotype combinations in host and pathogen populations. We consider that the total plant abundance is limited by a carrying capacity. For the pathogen, we track only genotype frequencies because the number of pathogen isolates to which a plant is exposed is low compared to the final pathogen density, such that relative pathogen frequencies determine host infection status.

The system is initialized from regional R-gene and effector pools. Motivated by the empirical distributions of effector and NLR repertoire sizes (FigureS1), we draw the size of the effector or NLR repertoire for each genotype at random from a Poisson distribution with mean *λ_E_* and *λ_R_*, respectively. The population is initialized with *n_p_*_0_ pathogen genotypes and *n_h_*_0_ host genotypes, constructed by uniformly sampling random combinations from equally sized effector and R-gene pools. As a simplifying assumption, we project all effector-NLR interactions, including indirect recognition mediated by intermediate plant proteins, to a bipartite effector-NLR recognition network. We assume that one effector can be recognized by exactly one R-gene, and vice versa. Susceptibility only happens in the presence of an effector and the absence of the corresponding R-gene. We later discuss possible extensions of the recognition network.

In each generation, seedlings germinate from the soil and each seedling encounters a random collection of pathogens. The number of pathogen strains each seedling encounters is drawn from a Poisson distribution with mean *β*. The genotypes of the infecting pathogens are sampled by multinomial sampling according to their frequencies. Note that this sampling scheme allows for stochastic extinction of pathogen genotypes whenever a pathogen genotype is not sampled in any infection. Additional host and pathogen genotypes are introduced by immigration with Poisson distributed rates *m_p_* (for pathogens) and *m_h_* (for hosts).

The fitness of a pathogen strain reflects its relative rate of growth across all hosts in one host generation. It is determined by two multiplicative factors that represent the detrimental effect of host recognition and the beneficial effect of unrecognized virulence factors. That is, a pathogen benefits from any of its effectors that evade host recognition and act as virulence factors to improve its ability to infect and reproduce, but is impaired by its effectors that are recognized and thus trigger host defenses. Plant fitness is determined by the infection outcome and the metabolic costs associated with expressing R-genes [17]. Failure to recognize pathogen effectors leads to plant disease, which reduces seed production.

For host *j* colonized by pathogens *i* = 1, 2*, …, n*, we define the fitness of each infecting pathogen strain *i*(*f_i_*) and the host fitness (*f_j_*) as follows:

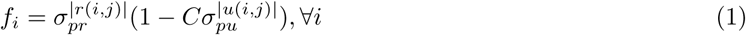

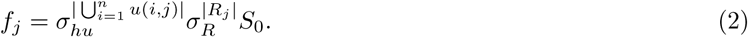

Here *r*(*i, j*) denotes the set of effectors that pathogen *i* possesses and are recognized by the host *j*, and *u*(*i, j*) denotes the unrecognized counterpart. Metabolic costs associated with R-genes are defined by the parameter *σ_R_* and the total number of R-genes *|R_j_|*. The selection strengths related to infection outcomes are specified by *σ_pr_* for the pathogen fitness discount due to recognition, *σ_pu_*for pathogen fitness gain due to escaping recognition, and *σ_hu_* for the host fitness discount due to recognition failure. We define a pathogen baseline fitness in the absence of virulence as (1 — *C*). That is, in the absence of either unrecognized effectors or recognized effectors (*|u*(*i, j*)| = *|r*(*i, j*)| = 0), the fitness of the pathogen genotype *i* is *f_i_* = (1 — *C*). The maximum number of seeds produced by a plant in one generation is defined by *S*_0_.

We simulate population dynamics for a grid of parameter regimes, defined by the strength of selection (*σ_pr_*, *σ_hu_*), fitness costs associated with the expression of R-genes 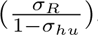, and the baseline fitness that a pathogen experiences when having no unrecognized effectors (1 — *C*).

The computational model generates different dynamics under different selection regimes. When plants experience strong selective pressure for defense success (small *σ_hu_*), often only one or a few well-adapted plant genotypes with relatively low numbers of R-genes dominate, and pathogens subsequently coexist neutrally due to relatively uniform host-imposed selective pressure (FigureS2 a, d). When pathogens experience strong selective pressure for evading host immunity (small *σ_pr_*) and hosts experience intermediate selective pressure (intermediate *σ_hu_*), genotype frequencies for both host and pathogen exhibit cyclic fluctuations, reflecting the advantage of being rare and the disadvantage of being common (FigureS2 b). The cycles are lost with increased *σ_hu_* due to slower dynamics among host genotypes (FigureS2 b, c). When host-imposed selection on pathogens is weakened (large *σ_pr_*), more recognized pathogen genotypes persist in the population (FigureS2 column e, f).

We then investigated the strain structure generated by the simulations to understand the processes maintaining strain coexistence. We focused on effector repertoire similarity, which reflects the strength of competition between strains for hosts, as well as their niche partitioning in terms of relying on different hosts for growth [77]. We addressed strain structure by constructing a strain similarity network and assessing its modularity. Each node in the network represents a strain, and edges represent pairwise effector repertoire similarity. The edge weights are defined by Pairwise Type Sharing (PTS) score, the number of effectors present in both pathogens (the cardinality of the intersection) over the total number of effectors of the two (see Methods 6.5), a quantity similar to the Sørensen index [90]. A network module represents a cluster of effector repertoires that have high overlap with each other but low overlap with repertoires outside the module (see Methods 6.6). The member strains of such a group would thus experience weaker competition for hosts with strains in other modules.

With a parameter sweep of *σ_pr_, σ_hu_,* and (1 — *C*) consisting of 50 replicates, we found that modular structures were generated when selection was strong for pathogens to evade immunity (*σ_pr_*≤ 0.5) and when selection was weak to intermediate for hosts to successfully recognize effectors (*σ_hu_* ≥ 0.5) (Figure1 a). These modular structures correspond to the co-diversification of the host and pathogen strains (Figure1 b). Temporal dynamics of the modules show that the modules are groups of pathogen strains that rely on the same hosts, and concurrently increase in frequencies (FigureS3). Because the simulation uses a large maximum plant seed production value (*S*_0_ = 10, 000) realistic for *A. thaliana*, the hosts often maintain a relatively constant maximum population density. Therefore, we also experimented with *S*_0_ = 100 and confirmed that our observations are robust to reduced maximum seed production (FigureS4).

**Figure 1:**
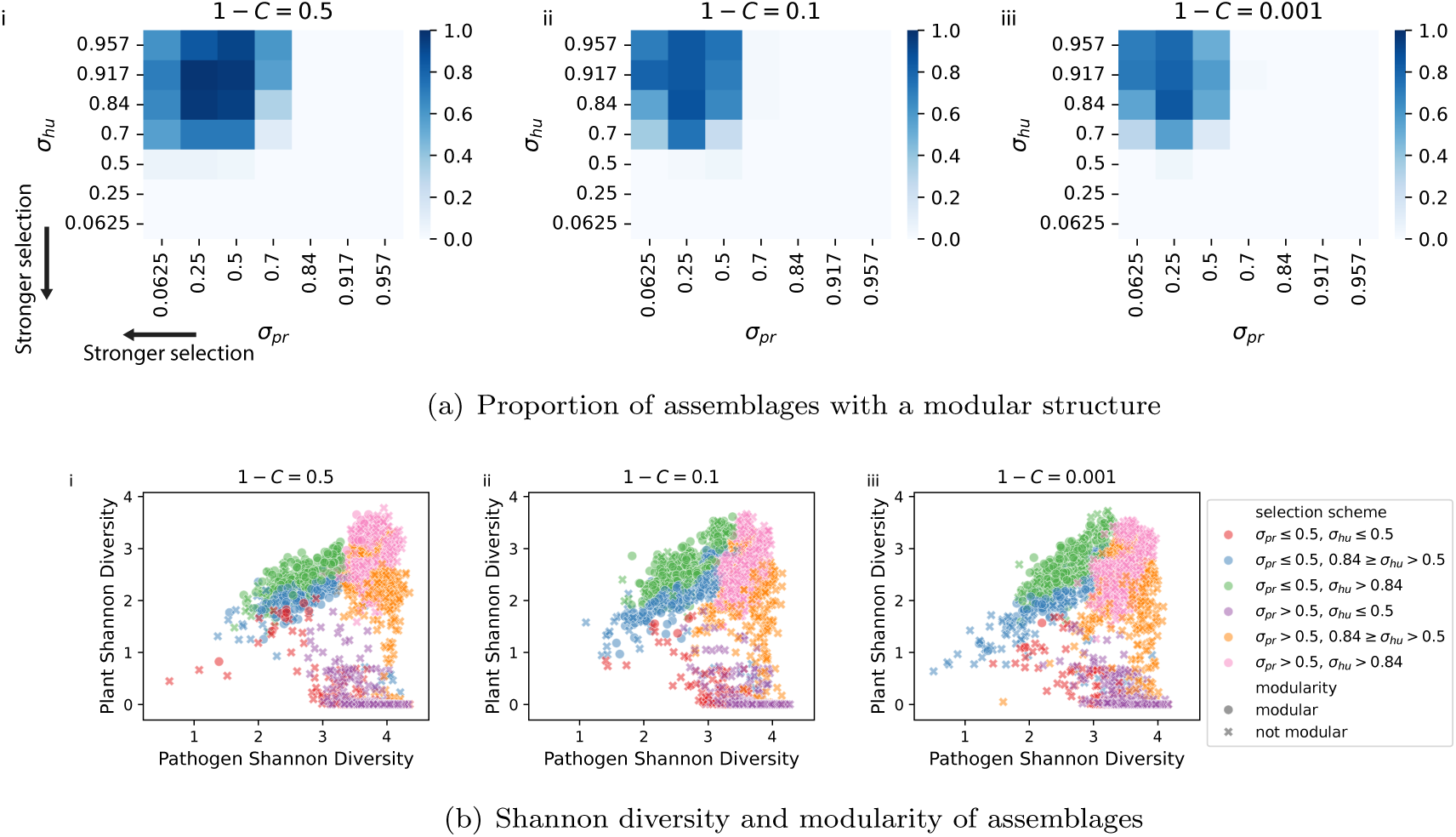
Modular structures are generated by the simulations, and are correlated with host-pathogen co-diversification. a) Modular structure is found in similarity networks defined by pairwise type sharing (PTS). The shade gradients represent the proportion of simulations (*n* = 50) with a modular PTS network structure after 1000 generations. b) Relationship between Shannon diversity and the structure of the resulting assemblages. Colors represent selection schemes defined by ranges of *σ_pr_* and *σ_pu_*: *σ_pr_* ≤ 0.5 and *σ_pr_ >* 0.5 for strong/weak selective pressure against the pathogen with recognized effectors; *σ_hu_* ≤ 0.5, 0.84 ≥ *σ_hu_ >* 0.5, *σ_hu_ >* 0.84 for strong/intermediate/weak selective pressure against the host when encountering unrecognized effectors. Whether the final assemblage has a modular similarity network is represented by a dot (modular) or a cross (not modular). For each realization, Shannon diversity is calculated by averaging over the last 200 generations to account for fluctuations. We used 50 realizations for each parameter combination of (1 — *C*), *σ_hu_*, and *σ_pr_*. Other parameters are: 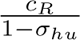 = 0.25*, β* = 3*, N_R_* = *N_E_* = 30*, n_p_*

### 2.2 *P.syringae* strains in nature exhibit effector similarity clusters that are phylogenetically aligned

For comparison with our simulation results, we consider a similar analysis of effector repertoires of *P. syringae* strains in nature. Although abundant *P. syringae* sequences are available from global sampling efforts, these are not suitable to address patterns of diversity in regional populations. Therefore, we sampled and sequenced 76 *P. syringae syringae* strains from the Midwestern US. These strains belong to *P. syringae* phylogroup 2, a widely present phylogroup that causes diseases in various plant species [16, 74]. Because effectors within the same homology family can have different virulence abilities [59], we analyzed effectors at the higher resolution of subfamilies [63] (although alleles within the same subfamily may still differ in virulence [23]). We then performed the same repertoire similarity analysis as on the simulation outputs. The constructed similarity network based on the PTS scores reveals a modular structure (*p <* 0.01, see Methods 6.6) (Figure2 a). The modules correspond to sets of *P. syringae* strains whose effector repertoires are more similar within than between the modules. Because our strains were isolated from four host species (*Arabidopsis thaliana*, *Cerastium vulgatum*, *Draba verna*, and *Lamium purpureum*), we further asked whether the modular structure reflects adaptation to host species. We find that the host of isolation does not correspond to repertoire module membership (Figure2 a).

**Figure 2:**
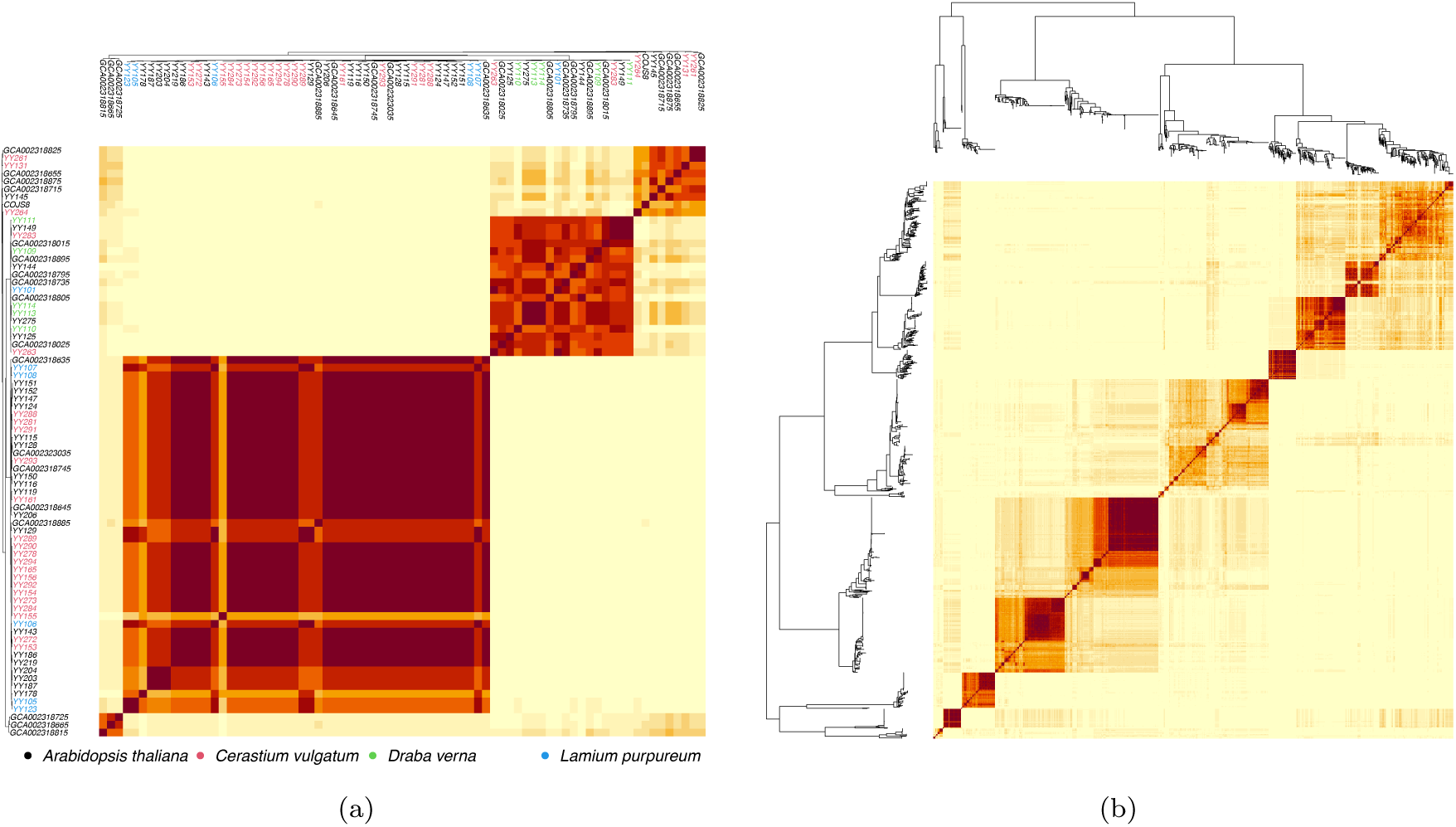
Pairwise type sharing (PTS) scores among P. syringae syringae strains from the Midwestern US (a) and the global dataset (b) show effector repertoire similarity clusters. The shade gradient represents PTS values from 0.0 (light; zero overlap) to 0.5 (dark; identical repertoires). In (a), strain names are colored by host of isolation.

To investigate the phylogenetic relatedness of effector repertoires among strains both between and within modules, we then built phylogenetic trees based on the core genomes. The modules of repertoire similarity exhibit a clear phylogenetic signal consistent with the core phylogeny (*p <* 0.01, see Methods 6.7), such that strains in different modules are associated with a higher phylogenetic divergence than strains within the same module. We address possible underlying processes in later sections.

Another pertinent observation is that the largest module corresponds to *P. syringae syringae* strains with only a few effectors and lacking a functional T3SS machinery [11], whereas strains outside this module carry at least 10 effectors (FigureS5). As this distinction may reflect different life history strategies that are outside the scope of our study, we restricted our downstream analyses to the strains with at least 10 effectors.

To investigate whether a pattern of modular structure of effector repertoires congruent with a phylogenetic signal is present in *P. syringae syringae* diversity at a global scale, we compiled existing datasets consisting of *P. syringae syringae* isolated from 37 countries and 98 host genera. These data allowed us to ask whether observed modularity maps to the host or location of isolation. For the former, we considered the six most frequent host of isolation genera in the dataset, *Actinidia*, *Prunus*, *Solanum*, *Arabidopsis*, *Coffea*, and *Phaseolus* (FigureS6). For the latter, we considered the most frequently sampled focal locations with subfamilies, which requires a higher sequence similarity than families, among *P. syringae syringae* strains. Among strains with at least 10 effectors, we identified 31 effector families, including 22 that occurred in more than one strain (Figure3). Multiple effector subfamilies are found within 11 of these effector families. It has been suggested that presence/absence patterns [31] of effector families can be attributed to gene loss and HGT [9]. In addition to these two mechanisms, the mosaic distribution of effector subfamilies may also be associated with recombination due to their high sequence similarity. Multiple effector subfamilies show distributional patterns consistent with genetic exchange. We further tested for the presence of recombination using the package PhiPack [22], and identified 13 effector subfamilies with mean refined incompatibility higher than random (Table 1) (see Methods 6.8). Our findings are consistent with previously reported evidence of effector recombination in *P. syringae* populations [33]. Thus, the modular structure of effector repertoires persists despite mixing from genetic exchange.

**Table 1:** Effector subfamilies identified in high-effector strains that exhibit a signal of recombination in PhiPack analysis.

| Effector | Estimated Diversity | Informative Sites | p-value |
| --- | --- | --- | --- |
| AvrE1k | 0.8 | 89 | 0.0 |
| AvrE1o | 1.5 | 61 | 0.0 |
| HopAA1o | 1.1 | 32 | 0.0 |
| HopB2d | 1.2 | 116 | 0.00002 |
| HopB2j | 0.8 | 116 | 0.0 |
| HopI1d | 2.8 | 59 | 0.0005 |
| HopI1l | 3.0 | 46 | 0.00041 |
| HopM-ShcM1g | 2.2 | 91 | 0.00011 |
| HopM-ShcM1x | 1.0 | 45 | 0.00018 |
| HopW1a | 0.8 | 60 | 0.0 |

**Figure 3:**
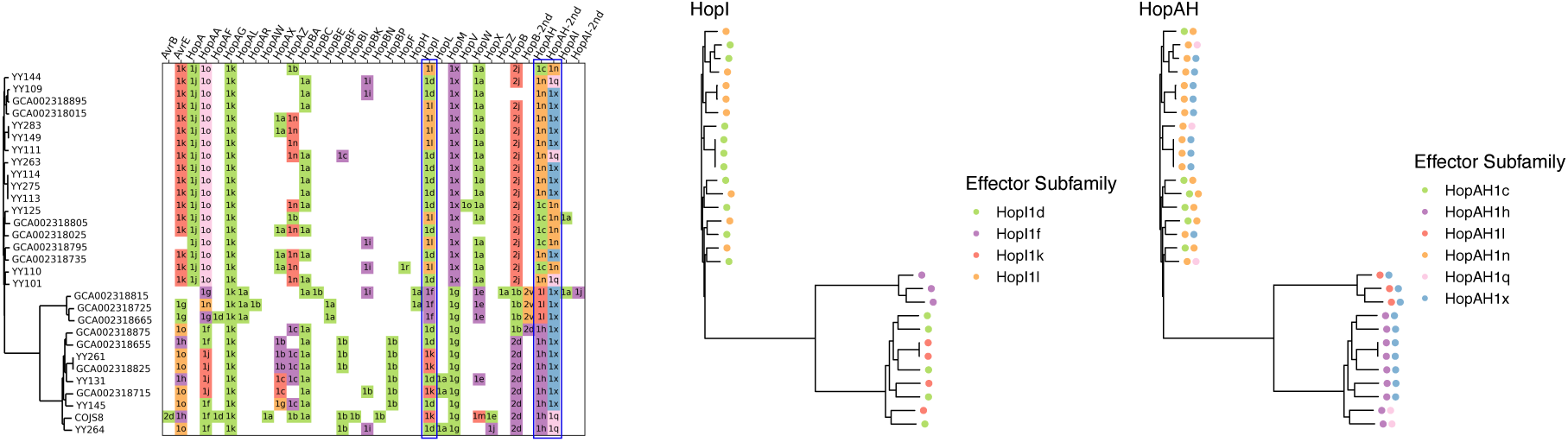
Effector subfamily distribution is consistent with patterns generated by genetic exchange. Left: effector subfamilies identified from high-effector strains. Each row represents a strain, ordered according to its core genome phylogeny. Each column represents one of the 31 identified effector families. Each colored square indicates the presence of an effector family (column) in a strain (row). Different subfamilies of the same family, as indicated by text, are marked by different colors in the same column. For strains containing more than one effector of the same family but different subfamilies, the two subfamilies are marked in separate columns (HopB and HopB_2nd, HopAH and HopAH_2nd, HopAI and HopAI_2nd). While some families have a subfamily distribution consistent with the phylogeny (ex. HopM and HopB), many exhibit patterns of mixing (ex. HopAZ, HopI, HopAH). Middle and right: zoom-in views of effector subfamily distributions for the HopAH (middle) and HopI (right) families. Dots of different colors represent subfamilies identified from the corresponding strain as indicated in the legend. When multiple subfamilies are identified for one strain, multiple dots are shown (right, HopAH).

### 2.4 NFDS from ecological interactions can contribute to the maintenance of phylogenetically aligned similarity clusters in the face of genetic exchange

Motivated by the evidence of both genetic exchange among subfamilies and a phylogenetic signal in the modularity of effector repertoires, we extended the model. Our original computational model is not suitable for examining pathogen population structure in the context of phylogenies, as it assembles initial and immigrant repertoires by randomly sampling from pools of effectors and R-genes. This setup implicitly allows for all possible repertoire combinations from their respective pools, and therefore mimics unconstrained and fast genetic exchange that would erase evolutionary history. To introduce phylogenetic relationships without modeling the full history that generated it, we start from a given population structure, superimposing a phylogeny, and ask about its maintenance rather than its origin. That is, we use an extension of the computational model to ask whether NFDS plays a role in maintaining population structure in the presence of mixing from effector recombination.

We address the role of NFDS in maintaining modules in the presence of genetic exchange by explicitly including phylogenetic structure and recombination events [39]. To this end, the communities are initialized with host and pathogen genotypes and their respective frequencies after running the original model for *t* = 100 generations. We assign a phylogeny to the strain structure by sampling from the distribution of pairwise strain phylogenetic distances of the high-effector strains from the Michigan *P. syringae* population. Our approach ensures that strains within an effector module have lower phylogenetic divergence on average than those outside of it (see Methods 6.9).

Recombination is modeled via random events in which an effector of one pathogen strain is uni-directionally replaced by that of another. In practice, this corresponds to effector replacement of one strain through homologous recombination [97]. We do not take horizontal gene transfer events into account, that is, the acquisition of additional effectors from another strain, because on the time-scale of interest, the rate of such events is lower than that of recombination [32, 31] and thus, is unlikely to alter the results qualitatively.

To represent biological barriers to recombination, we implemented two types of constraint ([80, 38]): (i) recombination between two strains becomes less frequent with increasing phylogenetic divergence [101, 66], and (ii) recombination between two gene segments becomes less frequent with increasing sequence divergence [89]. For (i) we define the probability of successful recombination as

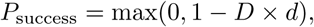

such that *D* is the phylogenetic distance between the two strains and *d* is a parameter controlling the rate of decrease in recombination frequency with increasing phylogenetic distance. For (ii), we restrict recombination events to effector pairs with sufficiently similar sequences. Following 2.2, we allow effectors in the model to represent effector subfamilies, and arbitrarily assign the original regional pool of effector subfamilies to *G* equal-sized families within which we consider effectors as sufficiently similar for recombination. In practice, the number of groups represents the strictness of this constraint, such that one single group implies that effector sequence differences have little effect on recombination frequency, whereas a large number of groups implies that only very similar effectors can recombine. We sample the number of recombination events at the end of every host generation from a Poisson distribution with mean *λ_recomb_* (see Methods 6.9). No immigration is allowed in the extended model.

We simulated the community under three different scenarios: (A) both pathogen recombination and NFDS from host-pathogen ecological interactions; (B) recombination only; (C) NFDS only. We analyzed the resulting assemblages at the end of 100 generations for modularity and phylogenetic signal. We selected seven initial conditions where strain structure is not lost in the NFDS only scenario of the model extension. For each initial condition, we ran 50 replicates each for scenario A, B, and C, with a range of recombination rates *λ_recomb_*. We evaluate the relative role of NFDS in promoting the persistence of strain structure (as defined by effector repertoires) by investigating the relative proportion of simulations where the modular structure is maintained after 100 generations under the joint action of NFDS and recombination compared to recombination alone for different combinations of *λ_recomb_*and *d*. We show that recombination without selection (scenario B) produces effector repertoires that erode preexisting modules, especially as rates of recombination increase (orange lines) (Figure4, S 8, S9). In contrast, NFDS contributes to the maintenance of the phylogenetically aligned modules in the presence of recombination (scenario A; blue lines). Phylogenetically closely related strains are not necessarily grouped into the same module, but the phylogenetic signal can persist (FigureS10 d). When the modular structure is preserved, the similarity modules are likely phylogenetically aligned (Figure4 d). Restriction of recombination due to phylogenetic distance (*d*) weakly stabilizes the modular structure in the absence of NFDS (Figure4 b, c, dotted orange lines). Assigning effectors to different numbers (*G*) of genetically similar recombining effector families has little effect at low *G*, but may contribute to the perseverance of phylogenetically aligned modules at high *G* (FigureS11) by strengthening differences between modules among which recombination rarely happens.

**Figure 4:**
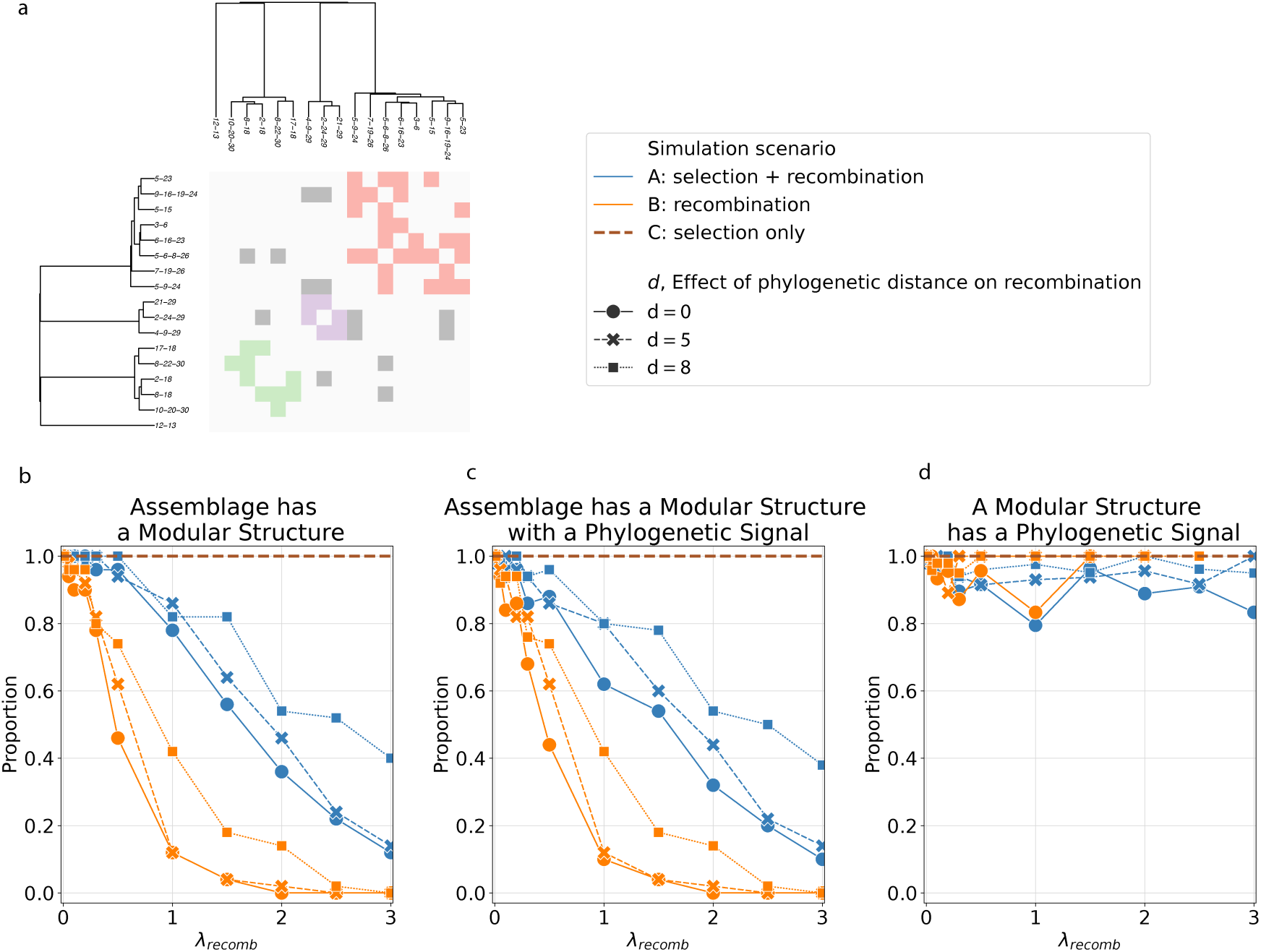
Maintenance of phylogenetically aligned clusters by NFDS in the presence of genetic exchange. (a) Initial condition of the simulations. Results on simulations with other initial conditions are reported in FigureS 8 and S 9. In (b-d), lines represent the proportion of assemblages obtained at the end of the simulation (g=100) with a modular structure (b), a modular structure with a phylogenetic signal (c), and with a phylogenetic signal given the structure is modular (d, calculated as (c) divided by (b)), under different modeling scenarios indicated in the legend above. Genetic exchange is allowed between any pair of effectors (*G* = 1). Constraints of recombination due to strain phylogenetic divergence, *d*, are marked by different line types. The initial assemblage contains 17 pathogen genotypes and 32 plant genotypes. Parameters: σ_pr_ = 0.25, σ_hu_ = 0.7, 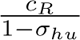 = 0.25, 1 – C = 0.9, β = 3, K = 20000, S_0_ = 10000.

The degree to which NFDS can maintain strain structure against the mixing by recombination varies based on initial conditions (FigureS8, S9, S12, S10, S13), although the general trend is consistent. We later discuss potential causes of this variation. We note that our simulations were run for 100 years from the constructed initial conditions. Had we used a shorter time, we would have seen a higher fraction of runs in which modules persist for a given recombination rate (FigureS12), thus illustrating the inter-dependence between the recombination rate and the speed of the eco-evolutionary dynamics of the assemblages.

### 2.5 The patterns of effector co-occurrence are not consistent with functional or physical linkage

Beyond NFDS, another potential mechanism for creating groups of co-occurring effectors, and hence modules of similar effector repertoires, is functional and/or physical linkage [67]. (Functional linkage refers here to pairs or sets of effectors that are not necessarily physically linked and result in the increased fitness of a strain carrying these). We analyzed co-occurrence patterns in effector repertoires across different locations at the level of provinces (see Methods 6.10) and identified clusters of similar co-occurrence patterns for each effector (FigureS15, S16 a, b). Since physical proximity between genes may increase their co-occurrence due to linkage, we calculated distances between effectors using all strains with complete assembly and repeated the analysis on this subset of strains. Representative results are illustrated in FigureS15. We found that most effectors co-occur with different sets of effectors across different groups of locations, forming clusters of co-occurrence patterns (FigureS16 a, c). For instance (FigureS15), three clusters are identified for AvrRpm1a, and more than three for HopAG1k. Using HopAG1k as an example, we also show that multiple clusters can be found for strains of the same phylogroup (phylogroup 2) isolated from a single host genus (*Prunus*). Locations that are close together may share co-occurrence clusters, yet some patterns are shared across distantly located provinces (FigureS15). In FigureS16 b, d), we summarize the proportion of all the effectors belonging to the same co-occurrence cluster for each pair of provinces (ordered by geographical proximity). Additionally, the majority of effectors are *>* 10^5^ bps apart (FigureS16 e), and therefore are not on the same pathogenicity island [2] or tightly linked. Thus, simple pairwise (or group-wise) functional or physical linkages do not provide a compelling explanation for the modules in effectors’ sharing between genomes, as one would expect more consistent associations across different locations.

## 3 Discussion

The growing characterization of the *P. syringae* pangenome offers an opportunity to understand the eco-evolutionary processes underlying strain diversity and coexistence [33]. Our work specifically addresses local *P. syringae* population structure in terms of the effectors repertoires in different genomes, and reports phylogenetically aligned similarity clusters. In contrast to previously proposed lifestyle differences, such as adaptation to agricultural versus wild hosts [51], we discovered modules among strains isolated from wild hosts that do not reflect host species nor location of isolation. We also detected recombination at a regional scale, suggesting that genetic exchange may erode modular patterns. Although the similarity clusters in the data are consistent with the modular population structure generated by NFDS in our initial eco-evolutionary model, our initial model was not sufficient to understand how phylogenetically aligned modules can be maintained. We therefore extended the model to account for both the phylogenetic signal of clusters and recombination, and showed that NFDS can counterbalance the mixing of effector repertoires by recombination and, in doing so, maintain strain structure.

We propose that NFDS can play a role in structuring and maintaining high *P. syringae* intraspecific strain diversity. With strong selective pressure against recognized effectors, our computational model produces modular population structures. This result is analogous to results obtained in other contexts. In ecological models with a one-dimensional trait axis, competition between species as a function of distance can drive the evolutionary emergence of coexisting groups of similar species [87, 29]. This partial form of limiting similarity enhances coexistence by lengthening persistence in sufficiently similar species while also reducing competition among those sufficiently different. Another example is the modular strain structure driven by competition for hosts through cross-immunity as a function of antigenic similarity in models of human host-pathogen systems [103, 45, 78] (e.g. coexisting malaria strains have weakly overlapping *var* repertoires [4]). The organization of pathogen genotypes into modules reflects niches emerging from ecological interaction and competition via immune memory, in contrast to preexisting niches such as lifestyle and habitats. For *P. syringae*, we can consider the niche of a module of strains as the set of host genotypes that they can infect. In our simulations, despite changes in module memberships due to immigration and extinction, the modules remain groups of pathogen strains that increase in frequencies concurrently by exploiting the same host genotypes (FigureS3 **e**, **f**). In subsequent generations, the exploitation of abundant hosts provides a fitness advantage to rare hosts and a subsequent rise in their frequencies. This in turn favors pathogen strains being part of other effector modules to rise in frequency. Therefore, modules, or coalitions of modules, partition hosts by both defense abilities and temporal frequency changes.

The dynamically generated niches we propose go beyond the current conception of *P. syringae* ecotypes, which is largely based on established niches of different habitats/life-history or host species (pathovars). [9, 33, 71]. The ecotype model describes lineage formation due to sweeps purging diversity among closely related strains, and reduced recombination among distantly related ones [26, 35, 40, 100]. In our model extension, we show that despite constrained genetic exchange across phylogenetic distances, NFDS is still required to counteract the erosion of strain structure by recombination. Although this requirement would be lifted when rates of recombination are very low or strongly constrained among closely related strains, this contradicts the observed mosaic distribution of effector subfamilies in the regional population. Moreover, the empirically observed depression in recombination rate across distantly related strains could also be promoted by selection against hybrid repertoires that fall outside the emergent niches of hosts R-gene repertoires. One subtlety is that some subfamilies are more narrowly distributed among closely related strains, including AvrE1k, HopA1j, HopAA1o, HopB2j, HopBP1b, HopM1x, HopM1g, and HopW1A, whereas others show extensive mixing, especially HopAH1x and HopI1d (Figure3 left). Interestingly, some of the narrowly distributed subfamilies also show evidence of recombination (Table 1). As there is currently no evidence of differences in recombination rates among effector subfamilies, this variation among distributional patterns may be attributed to transient dynamics, functional redundancy [23], or the persistence of repertoires overlapping with repertoires from more than one module, as observed in the simulation (FigureS3 d grey squares).

The generation and maintenance of niches by NFDS against introduced recombinants are robust on ecological time scales and under different initial strain compositions. In addition to selecting against recombinants recognized by hosts of both parents, in FigureS10, we show that strain structure reorganizes in response to perturbation by recombinant strains. The phylogenetically aligned modules could be eroded eventually with a high number of recombinant strains, and the ability to withstand this effect is stronger for some initial conditions compared to others (i.e. more robust with IC1 than with IC2, FigureS12). For example, in FigureS13, following the extinction of R-gene #11, the corresponding effector #11 becomes a prevalent member of the repertoires of many recombinant strains, and gradually erodes the strain structure. Restriction of recombination due to effector sequence similarity (high *G*) also promotes the robustness of strain structure (FigureS 11) because recombination with limited possibilities reinforces modules.

An explanation for the modular structure, alternative to niches from selection, is that of “linked effectors” that co-evolve, due to physical or functional linkage. We have presented evidence against it, with multiple patterns of co-occurring effectors that are also inconsistent across geography. While epistatic effects could reinforce and contribute to modularity, they do not appear as a dominant process.

A factor contributing to the uncertainty of our conclusions is the lack of report on the absolute rate of *P. syringae* recombination events in the literature. However, bacterial recombination is detected at ecological time scales [64]. We have experimented with a range of *λ_recomb_* values and found that the effect of NFDS in maintaining strain structure is most pronounced at intermediate *λ_recomb_* values, such that recombination is frequent enough to erode strain structure, while selection has enough time to respond.

It is worth noting that the setup of the extended model does not allow the dynamic re-organization of modules via the immigration of effectors or R-genes, including novel ones. This setup permitted us to remain agnostic as to how phylogenetically aligned models were generated in the first place. This question would require explicit evolution including innovation to track the relationship of the core genome to effector genotypes. We have implemented a more tractable approach, leaving further extensions of the computational model for future work.

Our work suggests a number of additional future directions. To further examine the effect of NFDS on strain coexistence, the dynamics of *P. syringae* populations could be studied over time by longitudinal yearly sampling of microbial strains at a local scale. Although considerably more costly, analyzing the distributions of NLR-repertoires of hosts (in terms of susceptibility to certain effector repertoires) would enable interrogating whether emergent niches arising from competition for susceptible hosts can be interpreted as a function of host genotype availability.

The current version of the computational model can be extended in various ways to gain further understanding of the eco-evolutionary dynamics of local pathogen strain structure. We currently use a recognition network structure which is a simplified version of the NLR-“hub”-effector tripartite interaction network found in some plant host-pathogen interactions, in which the “hubs” are plant proteins that mediate NLR-effector recognition (for example, guardees) [72, 50]. Given the presence of this intermediate layer, it is natural to move beyond a one-to-one recognition network (e.g. an asymmetric recognition structure [52]). Understanding the effect of recognition network structure on eco-evolutionary dynamics will contribute to explaining observed diversity and strain structure patterns. Moreover, it has been shown that the effector repertoire composition is not the sole determinant of virulence [83], probably due to complex molecular cross-talks [72, 41]. Further extension of the computational model with an effector interaction matrix may elucidate some effector co-occurrence and avoidance patterns. A future model extension could also include linkages between effectors [69], as well as between effectors and the core genome, to capture the dynamical outcome of selection on effector repertoires.

An intriguing aspect of *P. syringae* populations is the extent to which strain coexistence is mediated by the accessory genome. On the one hand, accessory genome evolution may be restricted by the evolutionary history of a species (“phylogenetic inertia”) [65]; while at the other extreme, partially due to their high mobility, the accessory genome may also evolve independently of the core genome [14]. Although our work suggests that the *P. syringae* accessory genome is tightly correlated with the core phylogeny, we propose that instead of passively tracking core genome evolution, it may actively participate in structuring strain diversity. This is the case in another microbe in which NFDS is proposed as an important mechanism maintaining accessory genome diversity, *Streptococcus pneumoniae* [28, 43], where a model based on the frequencies of three thousand loci of the accessory genome and negative frequency-dependent selection successfully predicted the trajectories of strain prevalence after a major perturbation from vaccination targeting a subset of serotypes [7]. Thus, NFDS experienced by the accessory genome might be a common force shaping microbial evolution.

## 4 Data availability

All code for the computational model and analyzing strain structures is available at pascualgroup/ndfs_recomb_pseudomonas_arabidopsis. All code and data for gene prediction, and a table of all genomes including metadata are available at code_EL. The repositories will be made publicly available upon submission. The assemblies of all original sequences can be found associated with NCBI BioProject PRJNA1195362.

## 5 Funding and Acknowledgment

This work is a contribution of the GEMS Biology Integration Institute, funded by the National Science Foundation DBI 422 Biology Integration Institutes Program, award no. 2022049. This work is also funded by support from the Simons Foundation to JB. HM received support from the Center for Genomics and Systems Biology, NYU Abu Dhabi. The authors are grateful for the generous support of the Zegar Family Foundation, and would like to thank the staff of the Genomics Core at the NYU Center for Genomics and Systems Biology. The authors also thank Choghag Demirjian, Frédéric Labbé, and Hannah Whitehurst for valuable discussions to improve the manuscript.

## 6 Methods

### 6.1 Strain isolation and sequencing

*P. syringae* strains sequenced in this study were either isolated from *Arabidopsis thaliana* plants collected from Michigan in 2022 or from *Arabidopsis thaliana*, *Draba verna*, *Cerastium vulgatum*, or *Lamium purperum* plants collected from Michigan and Indiana in 2003-2005 (see data repository 4). Ground plant material was streaked onto either Kings B media with nitrofurantoin or R2A. Colonies with the morphology of *P. syringae* were taken for validation by PCR performed either on the *gyrB* or *shc* genes to identify *P. syringae* [12]. Isolates confirmed as *P. syringae* were grown in LB, and DNA for sequencing was extracted using a SPRI-bead based method. Libraries were prepared with the Nextera XT DNA Library Prep Kit and sequenced on an Illumnia NextSeq with a 2×75 read configuration.

### 6.2 Genome assembly and annotation

Genomes were assembled with SPAdes (v 4.0.0) using default settings [10]. Prodigal (v 2.6.3) and Augustus (v 3.4.0) were used to annotate coding regions. All coding regions that were called complete by at least one program were included in the predicted proteome for a given genome [46, 92]. We combined the results of the two gene prediction programs by parsing their GFF results using a custom Python script. Identical gene calls from both programs were filtered so that only one remained. When the programs predicted different genes in the same position, the longer gene was taken for subsequent analysis. Effectors were identified by BLASTing (blast+ v 2.13.0) the predicted proteome (amino acid sequences) of each genome against a custom database consisting of a single representative of each subfamily in the PsyTEC collection [3, 59]. Proteins that had a hit in the database with at least 80% identity, an E-value of less than 1*e^—^*^5^, and an alignment that covered at least 80% of both the query and target sequence were considered effectors. Putative effectors were assigned to the subfamily to which they had the match that passed the previous criteria with the highest bit-score. Core genome phylogenies were constructed using panX (v1.5.1) with the core gene threshold set to 95% (-cg 0.95) and otherwise default parameters [34].

### 6.3 Compiling *P. syringae* global set

To supplement the collection of *P. syringae* genomes we sequenced from the Midwestern US, we collected genomes with species names consistent with being members of the *P. syringae* species complex from NCBI and IMG. The full list of species names we considered is listed in the data repository 4. We also included all genomes from the BioProjects PRJEB24450 and PRJNA292453, [49, 32] as both projects represented large-scale efforts to sequence *Pseudomonas* plant pathogens.

For all genomes we then used fastANI (v 1.33) run with default settings [47] to separate members of *P. syringae* from the closely related species *Pseudomonas viridiflava.* We excluded *P. viridiflava* isolates because they tend to have few effectors, and are necrotrophic. *P. syringae* is hemibiotrophic, and thus is likely to interact with the host immune system differently[49]. If a genome had a higher mean average nucleotide identity (ANI) to a set of known *P.* syringae genomes than to a set of known *P. viridiflava* genomes, we considered the genome to be a *P. syringae* genome regardless of the species identifier. We provide the set of known P. syringae and *P. viridiflava* genomes in 4. We chose the 11 *P. viridiflava* genomes deposited on NCBI which are complete at at least the chromosome level. For *P. syringae* we chose assemblies for two strains from each of the primary phylogroups from [32]. For phylogroup 2 we chose two strains that were sequenced in [50].

To remove duplicated genome assemblies likely due to sequencing different isolates of the same strain, we ran FastANI on all the identified *P. syringae* genomes, and kept one sequence for all pairs of genomes with an ANI of 100%.

### 6.4 Eco-evolutionary model

#### Genotype initialization

The regional pools of effectors and R-genes were defined by *N_E_*= 30 unique effectors and *N_R_* = 30 unique R-genes. Each R-gene recognizes exactly one effector. For each realization of the simulation, we started our simulation with *n_p_*_0_ = *n_h_*_0_ = 20 unique pathogen/host genotypes. Each genotype was constructed as a random collection of effectors/R-genes from the regional pool. We first drew the number of effectors/R-genes for the genotype from a truncated Poisson distribution (≤ 30 effectors/R-genes), with mean *λ_E_* = 6 for effectors and mean *λ_R_* = 12 for R-genes. The identity of effectors/R-genes were then drawn without replacement from the local effector/R-gene pool. After all initial pathogen/host genotypes were constructed, their initial frequencies were drawn from a uniform distribution **U**[0,1] and then normalized to obtain a sum of 1. Initial host genotype frequencies were converted to initial genotype densities by multiplying the initial seed density *S*_0_ = 10^5^ by the respective genotype frequency.

#### Seed germination and seedling survival

In each generation, all seeds germinate with a probability *p_germination_*. We calculated the number of surviving seedlings (*N_survival_*) as a function of the total number of germinating seedlings (*N_total_*) and the carrying capacity (*K*) using a Beverton-Holt recruitment function

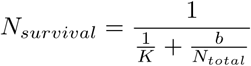

[18], and each seedling was then sampled for survival with a probability *<u>^Nsurvival^</u>*. Because *p_germination_* and *b* (at the limit of large *K*) can be absorbed by rescaling the maximum seed production *S*_0_ in computation, we used *p_germination_*= 1, *b* = 1 for all simulations to reduce the number of parameters with which we experimented.

#### Host and pathogen abundance/frequency update

In each generation, the infection outcome of each plant determines the fitness of the host and its associated pathogens as defined in Equation 1, 2. The range of each parameter can be found in Table 2. We calculated the relative fitness of each pathogen strain as its sum of fitness on all hosts divided by the sum of the fitness of all pathogen strains on all hosts. We normalized the relative fitness of all pathogen strains to obtain strain frequencies for the following generation.

**Table 2:** Description of fitness parameters in Equations 1, 2.

| Parameter | Definition | Range |
| --- | --- | --- |
| $\sigma_{pr}$ | Fitness discount on pathogen due to a recognized effector | [0,1] |
| $\sigma_{pu}$ | Discount on $C$ (see the last row) by an effector that successfully exploits the host | [0,1] |
| $\sigma_{hu}$ | Fitness discount on host due to an unrecognized effector | [0,1] |
| $\sigma_R$ | Fitness discount on host due to metabolic cost of an R-gene | [0,1] |
| $C$ | Fitness penalty of unable to exploit the host | [0,1] |

The seeds produced by each plant as in Equation germination process described above.

### 6.5 Pairwise type sharing

were added to the pool of seeds, and subjected to the We calculated pairwise type sharing (PTS) values to measure the similarity of effector repertoires between two strains *S_i_* and *S_j_*. For strains *S_i_* and *S_j_*, PTS is defined as,

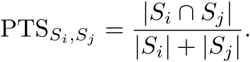

where *|S_i_* ∩ *S_j_|* is the number of effectors shared by both strains and *|S_i_|* (*|S_j_|*) is the total number of effectors in strain *i* (*j*). We constructed a PTS network in which each node represents a single strain. For every pair of nodes representing strains *S_i_* and *S_j_* (*S_i_* /= *S_j_*), we added an undirected edge *E_Si_,S_j_* with weight PTS*_Si_,S_j_* if PTS*_Si_,S_j_ >* 0.

### 6.6 Network modularity

Network modularity was calculated using the R Infomapecology package (version 2.2.0) with default parameters [36]. Infomapecology is based on the Infomap algorithm, which partitions the network to minimize the information needed for describing the movement of a random walker on the network [82, 36]. For statistical tests of modularity, we used the following number of randomizations (*r*): *r* = 100 for the compiled global dataset, and *r* = 200 for both the Midwestern US population and populations generated by the computational model. We identified a network as modular if p-value *<* 0.05. In analyses of simulated assemblages without genetic exchange, we filtered for pathogen frequencies *>* 0.005 to reduce the impact of immigrant strains that went extinct quickly. We considered those low-frequency strains in the model extension because many strains may coexist at low frequencies when under high rates of recombination, especially when selection is removed.

### 6.7 Phylogenetic signal

The phylogenetic signal of effector repertoire modules was analyzed following the approach in Pilosof et al. [79] The original pairwise phylogenetic distance was calculated using the R Ape package (version 5.8) [76]. For each module, we then calculated *D*_obs_, the mean pairwise phylogenetic distance between all strain pairs in the module. We computed the average *D*_obs_ across all modules to obtain *D*_obs_. To generate a null distribution of mean phylogenetic distances within modules, we permuted the strains and obtained 100 shuffled networks that preserve module numbers and sizes. We calculated the same metric, *D*_shufie_, for all shuffled networks, and compared *D*_obs_ to the distribution of *D*_shufie_. A significantly smaller average observed pairwise distance compared to the shuffled pairwise distance (p-value *<* 0.05) indicates that strains within the same module are more closely related than expected by chance.

### 6.8 Identification of genetic exchange

We used the program PhiPack (Bioconda) [22] to identify recombination within effector subfamilies. For each effector subfamily present in the set of genomes from the Midwestern US, we constructed a DNA multiple sequence alignment of the coding sequences for all representatives of that subfamily with Mafft (v. 7.475) using the mafft algorithm with default settings [53]. We then ran PhiPack on the alignment using default parameters, except for increasing the number of permutations from 1000 to 100000 (-p 100000). All effector subfamilies with a Bonferroni corrected PhiP-value less than 0.05 were inferred to have undergone recombination.

### 6.9 Eco-evolutionary model with recombination

#### Initialization

We simulated the eco-evolutionary model as described previously for 1,000 generations to initialize our eco-evolutionary model with recombination. We first removed all genotypes with frequency *<* 0.005, consistent with the protocol used for modularity analyses, followed by renormalizing the frequencies. Next, we identified all effector repertoire modules in the pathogen strains (see Methods 6.6). We then assigned core genome phylogenetic distances based on an empirical distribution obtained from our strains sampled in Michigan. To do so, we calculated pairwise core genome phylogenetic distances of Michigan strains using the cophenetic.phylo function in R package Ape (version 5.8). The distances follow a bimodal distribution with two non-overlapping modes, corresponding to within- and between-module pairwise phylogenetic distances (FigureS14 left). We generated probability density functions for within- and between-module distances, respectively using the density function in R (with bw=“SJ”) (FigureS14 middle and right). We sampled within- and between-module pairwise distances for the pathogen strains from these distributions. We then constructed a UPGMA tree (R package phangorn, version 2.11.1) based on these pairwise distances and used the resulting tree to re-calculate phylogenetic distances by calling cophenetic.phylo. We repeated this process to generate multiple initial conditions. We required each initial condition to satisfy the following conditions: (1) the pathogens have a modular strain structure; (2) the modules exhibit a significant phylogenetic signal after assigning phylogenetic relationships; (3) after simulating the assemblage for 100 generations with selection and without recombination events (scenario C in 2.4), the strains exhibit phylogenetically aligned modules.

#### Recombination events

We simulated each recombination event by randomly picking a donor and a recipient strain with probabilities proportional to their frequencies. A donor effector was sampled uniformly from the donor strain’s effector repertoire. For the recipient strains, a recipient effector was sampled uniformly from all effectors in its effector repertoire that were different from the donor effector but belonged to the same family. The two strains were resampled if no such recipient effector exists to conserve the total number of recombination events. A new strain was created by mutating the recipient strain with the recipient effector replaced by the donor effector. The new strain has a phylogenetic distance of zero to the recipient strain, and the same phylogenetic distances to other strains as those of the recipient strain.

#### Rates of recombination events

At the end of each host generation, the number of recombination events was sampled from a Poisson distribution with mean *λ_recomb_*. We investigated values of *λ_recomb_*in the range [0.01, 3]. We determined this range based on a back-of-the-envelope estimation of the *P. syringae* effective recombination rate [97]. Derivation and relevant parameters are detailed in Appendix 7.1.

### 6.10 Effector co-occurrence patterns in the empirical dataset

#### Identifying patterns of effector linkages

For each focal effector subfamily *i* in the empirical dataset, we calculated a vector **v***^p^*of effector co-occurrences for each province (*p*). Each entry of **v***^p^*, **v***^p^*, represents the number of strains containing both *i* and *j*, divided by **v***^p^*. Here, a province represents a geographic location (i.e. an administrative region, for example, a state or a prefecture) from which strains in our global dataset have been sampled (see data repository 4). We then performed k-mean clustering on all the resulting vectors **v***^p^*’s of the focal effector *i* using Python scikit-learn package (v1.4.1.post1, with 2 to 15 clusters under default settings). We selected the best number of clusters, *k*, using the maximum Silhouette score. Additionally, we checked gap statistics of the clustering results [96], and classified the vectors as belonging to a single cluster if this choice yielded a higher gap statistic than multiple clusters.

#### Effector co-occurrence clusters shared across locations

For every pair of provinces, we counted the number of co-occurrence clusters for which both are classified together across all focal effectors, normalized by the number of effectors found in both provinces. A high (low) value indicates that these locations tend to share similar (distinct) patterns of effector co-occurrences For visualization, we ordered the provinces by their geographical distances. To do so, we first used the Python package geopy (v2.4.1) to obtain coordinates for all provinces with Nominatim service and to calculate all pairwise geodesic distances. We performed hierarchical clustering on the distances with the Python scipy package (v1.12.0) and used the resulting order of the leaves to order the provinces for visualizations in FigureS16.

