## Supplementary Materials for "Negative frequency-dependent selection contributes to modular structure of effector repertoires in *Pseudomonas syringae*"

### 7 Appendix

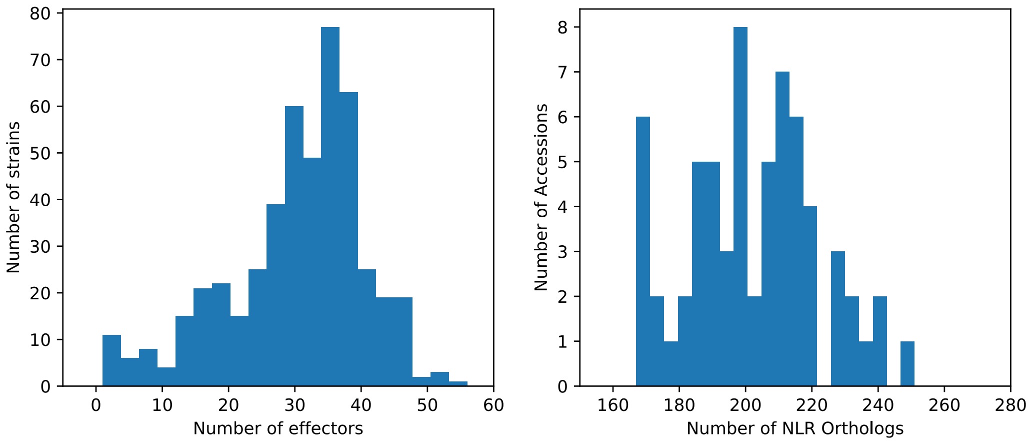


Supplementary Figure 1: Distribution of the number of effectors and NLRs from *P. syringae* strains and *A. thaliana*. Data are retrieved from published *P. syringae* accessions [32] and the *A.thaliana* pan-NLRome project (https://github.com/weigelworld/pan-nlrome) [99].

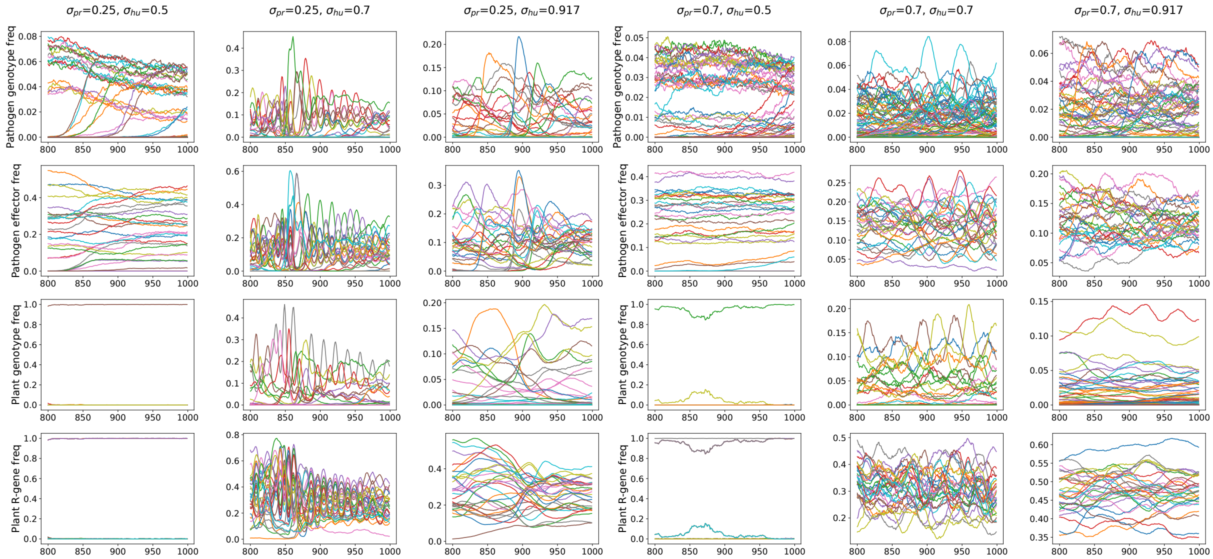


Supplementary Figure 2: Frequency trajectories of representative realizations under various parameter regimes (one realization for each regime). Only the last two hundred generations are shown. From the top to the bottom row, lines represent frequencies of pathogen genotypes, effectors, plant genotypes, and R-genes, respectively. Frequencies of effectors are calculated by pooling all pathogen genotype frequencies containing the effector, and therefore represent proportions of the population containing the effector. R-gene frequencies are calculated in the same manner. The colors are recycled. Parameters: $\frac{c_R}{1-\sigma_{hu}} = 0.25, (1-C) = 0.1, \beta = 3, N_R = N_E = 30, n_{p_0} = n_{h_0} = 20, K = 20,000, S_0 = 10,000, m_p = m_h = 0.5$.

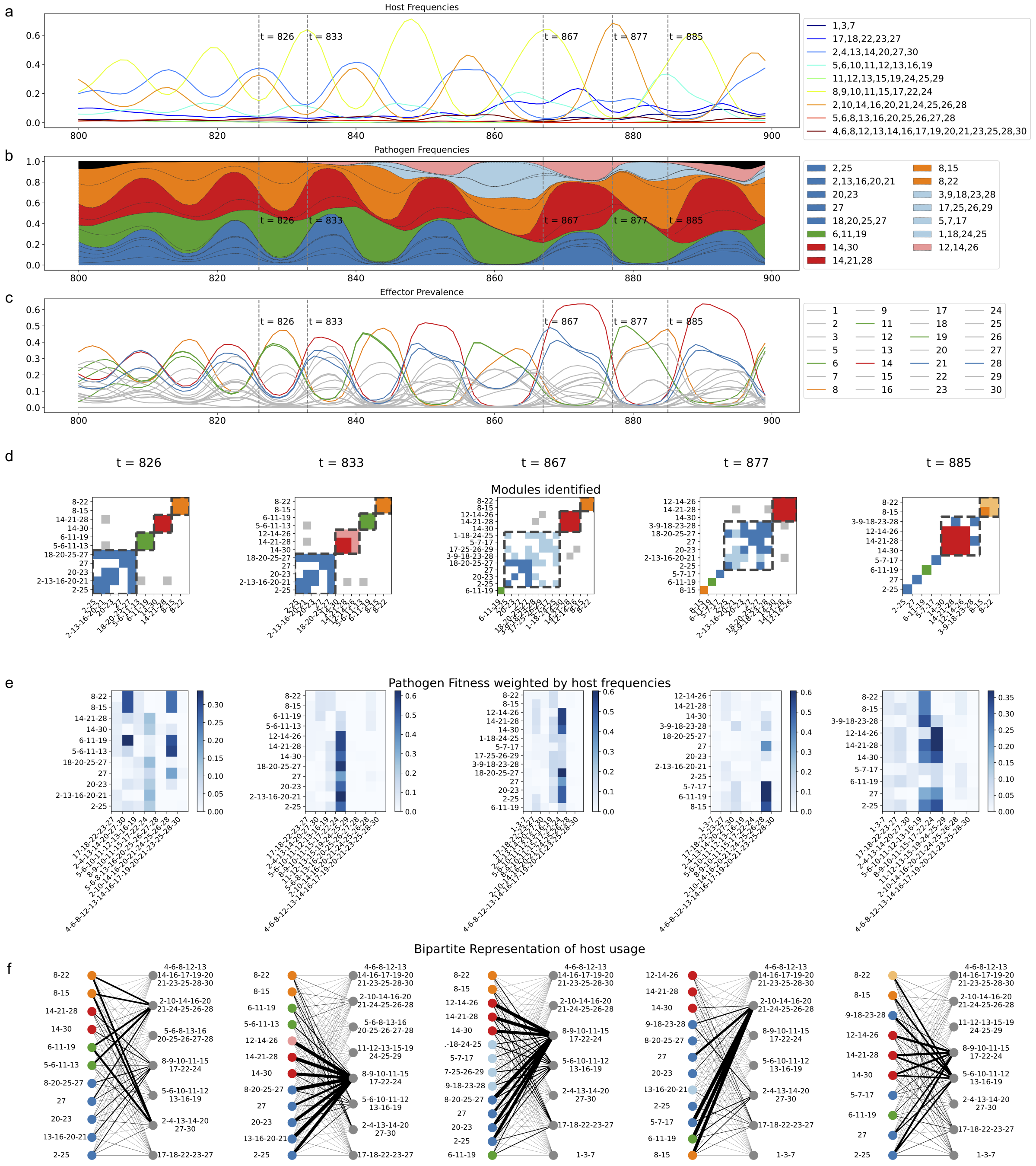

Supplementary Figure 3: Temporal dynamics of module memberships and host usages. a), b), c): Changes of host frequencies, pathogen frequencies, and effector prevalence (proportion of strains carrying a certain effector) through time. Pathogen frequencies (b) and effectors with maximum prevalence $> 0.4$ (c) are colored by the modules they are first identified as members, as in (d); black in (b) represents all strains with frequencies $< 0.005$. d) Modules identified at given generations are indicated with dashed boxes. The presence of an edge (PTS $> 0$) is indicated by a colored square, such that intra-module edges are colored by the first module the strain is identified with (at $t = 826$), and inter-module ones by grey. Light colors indicate new members of the modules. e) Shade represents the fitness of a pathogen (row) on a host (column), weighted by host frequency. f) The widths of the edges are proportional to the fitness in (e). The pathogen nodes are colored according to (d). Strains are filtered for frequencies $> 0.005$. Parameters: $\sigma_{pr} = 0.25, \sigma_{hu} = 0.7, \frac{c_R}{1-\sigma_{hu}} = 0.25, (1 - C) = 0.1, \beta = 3, K = 20,000, S_0 = 10,000$.

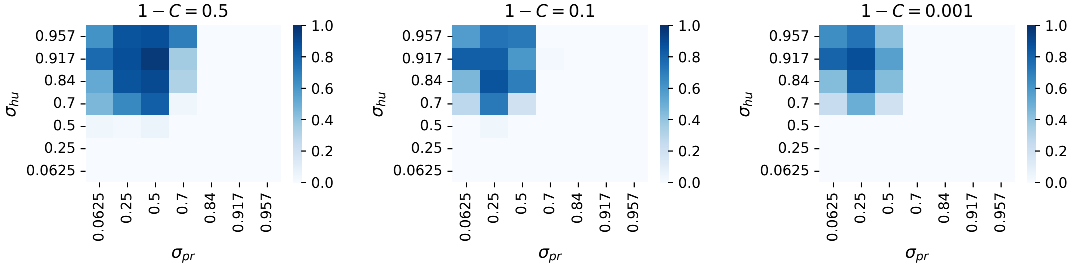


(a) Proportion of assemblages with a modular structure

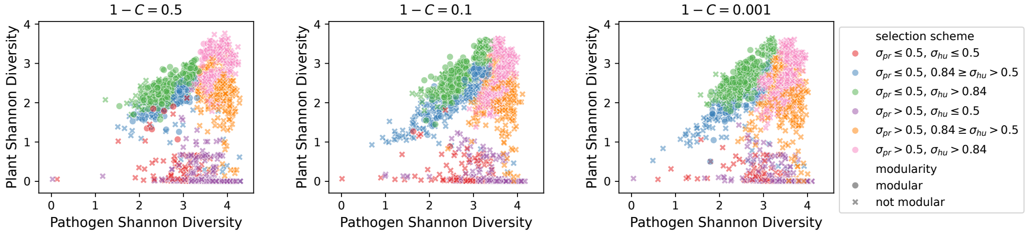


(b) Shannon diversity and modularity of assemblages

Supplementary Figure 4: Comparing to Figure 1, the observed pattern of the occurrence modularity structure is robust to a smaller basic reproductive rate $S_0$ (reduced to 100 from 10,000.). a) Modular pairwise type sharing (PTS) network structures were observed. The shade gradients represent the proportion of simulations ($n = 50$) with a modular PTS network structure after 1000 generations. b) Relationship between Shannon diversity and the structure of the resulting assemblages. 50 realizations are used for each parameter setting. Other parameters: $\frac{c_R}{1-\sigma_{hu}} = 0.25, \sigma_{hu} = 0.917, \beta = 3, N_R = N_E = 30, n_{p_0} = n_{h_0} = 30, K = 20,000, m_p = m_h = 0.5$.

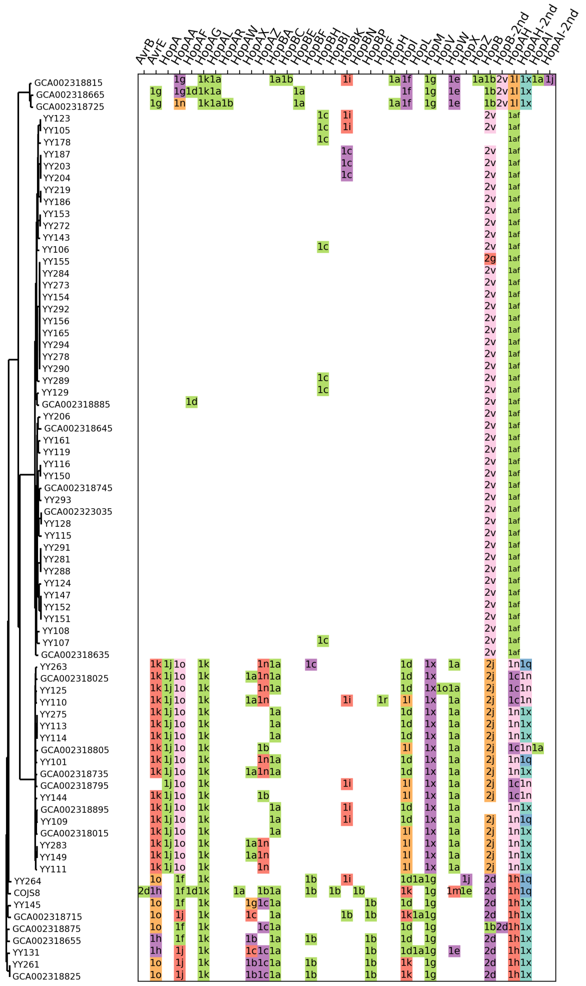


Supplementary Figure 5: Identified effector subfamilies from all Midwestern US strains. Similar to Figure 3, each colored square indicates the presence of an effector family (column) in a strain (row). Different subfamilies of the same family, as indicated in the text, are marked by different colors. For strains containing more than one subfamily of the same family, the two subfamilies are marked in separate columns (HopB, HopAH, HopAI). While some families have subfamily distribution consistent with the phylogeny (ex. HopM and HopB), many exhibit patterns of mixing (ex. HopAZ, HopI, HopAH).

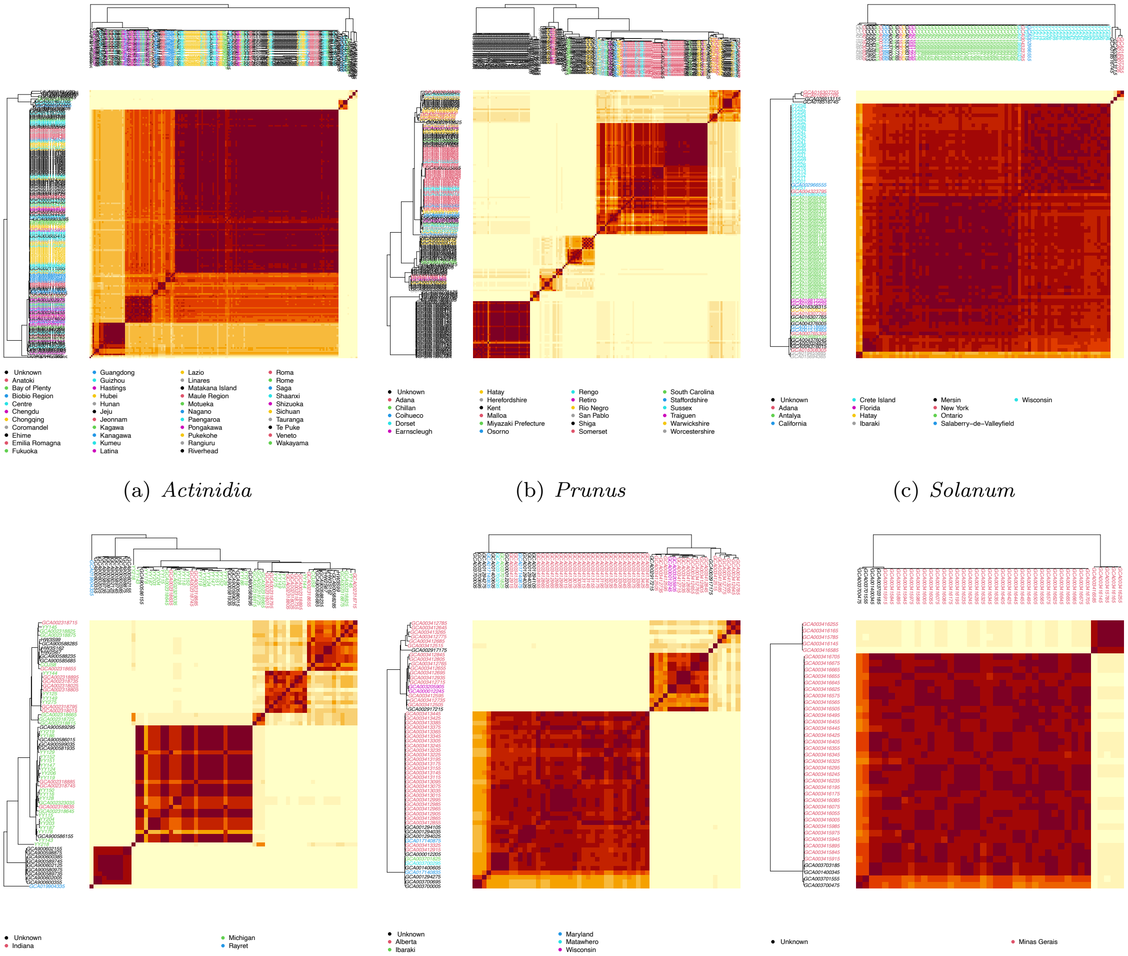


(a) *Actinidia* (b) *Prunus* (c) *Solanum*

(d) *Arabidopsis* (e) *Phaseolus* (f) *Coffea*

Supplementary Figure 6: Pairwise type sharing scores among *P. syringae* strains isolated from the six most abundant host genera in the global dataset. Colors represent locations of isolation [73]. Note the colors are recycled. The shade gradient represents PTS values from 0.0 (light) to 0.5 (dark).

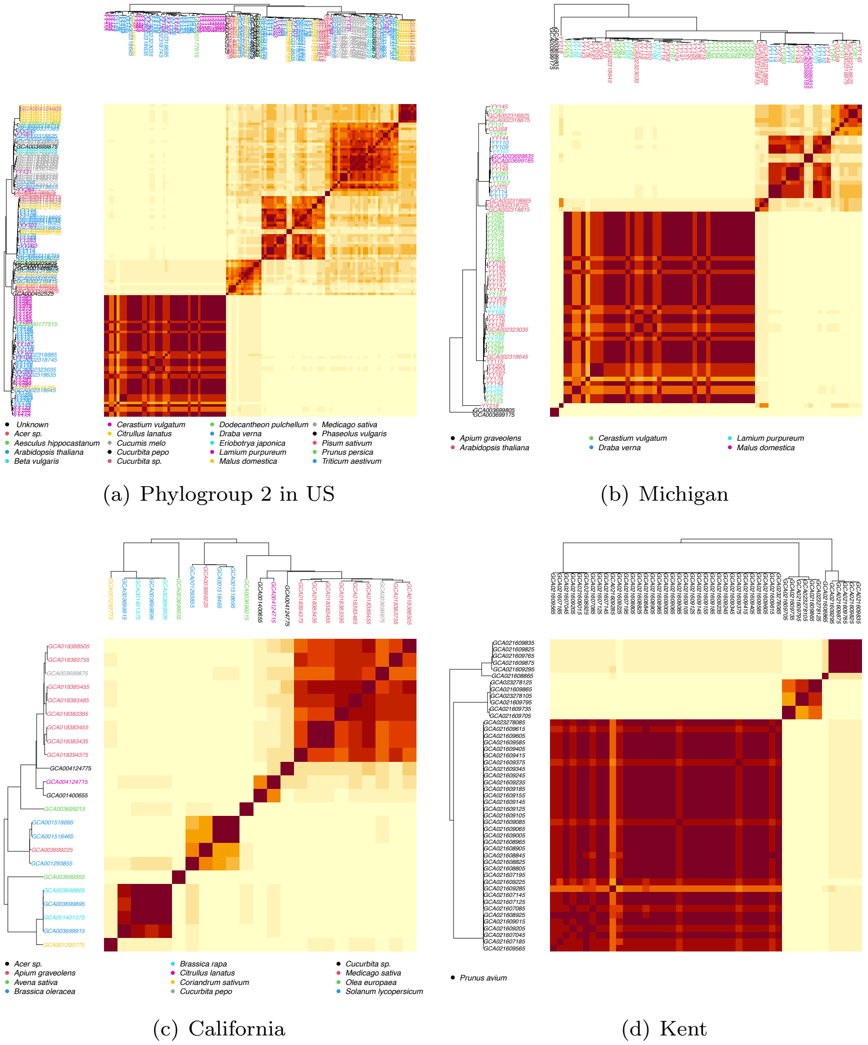


(a) Phylogroup 2 in US

(b) Michigan

(c) California

(d) Kent

Supplementary Figure 7: Pairwise type sharing scores among *P. syringae* strains isolated from four focal locations in the global dataset. Shown is PTS for (a) all phylogroup 2 strains collected from the US; (b), Michigan, (c), California, provinces where the highest number of strains are collected from more than five relatively evenly sampled host species; and (d), Kent, the province where the highest number of strains are collected from a single host species. Colors represent hosts of isolation [73]. Note the colors are recycled. The shade gradient represents PTS values from 0.0 (light) to 0.5 (dark).

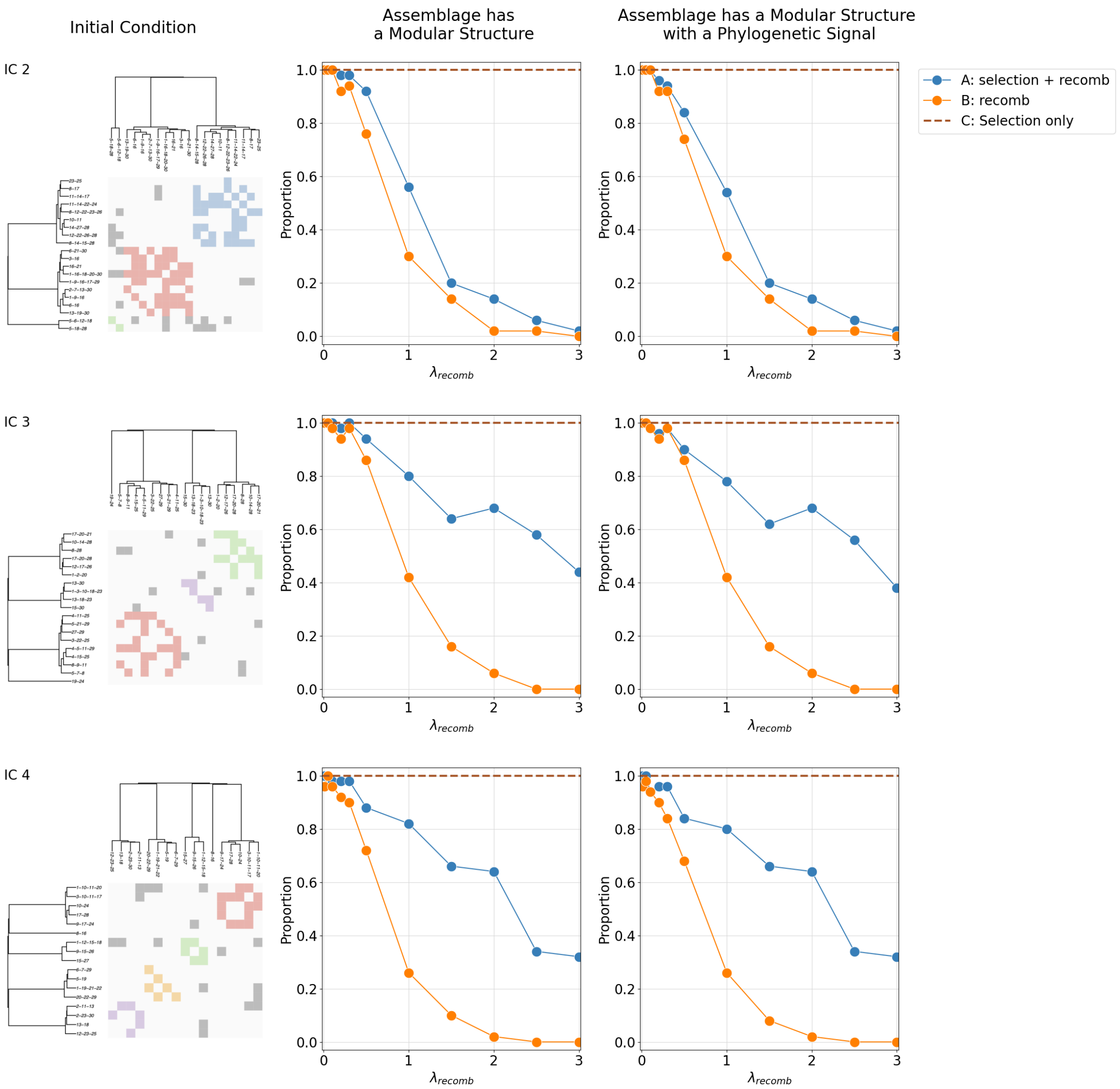


Supplementary Figure 8: The role of selection in maintaining phylogenetically aligned modules under different initial conditions. Initial conditions are shown in the left column and labeled by IC#s. Lines represent the proportion of assemblages obtained at the end of the simulation with a modular structure (middle) and a modular structure with a phylogenetic signal (right) under different scenarios, as shown in the legend. Parameters: $\sigma_{pr} = 0.25, \sigma_{hu} = 0.7, \frac{c_R}{1-\sigma_{hu}} = 0.25, (1 - C) = 0.1, \beta = 3, K = 20,000, S_0 = 10,000$

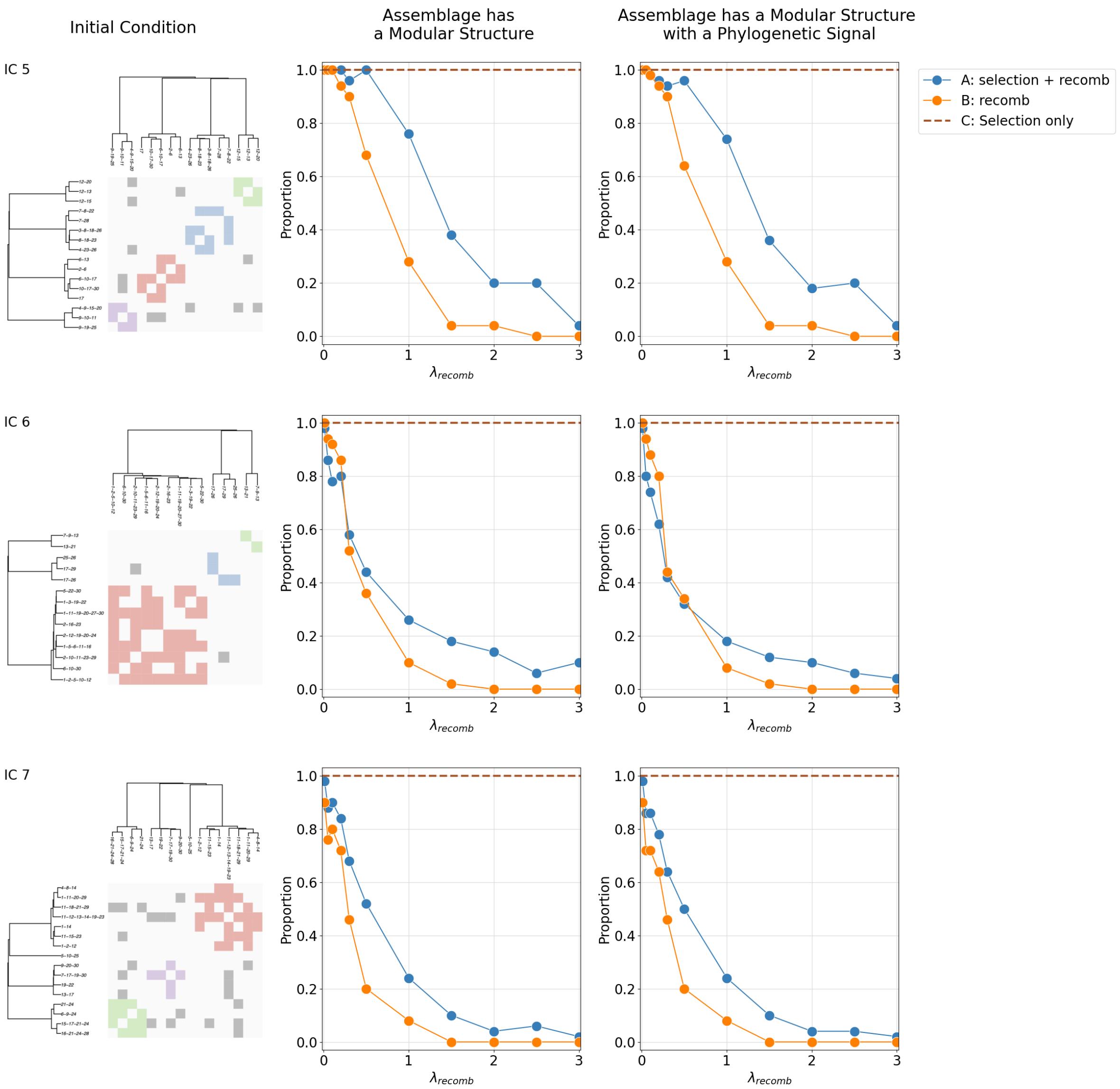


Supplementary Figure 9: The role of selection in maintaining phylogenetically aligned modules under different initial conditions (continued). Initial conditions are shown in the left column and labeled by IC#s. Lines represent the proportion of assemblages obtained at the end of the simulation with a modular structure (middle) and a modular structure with a phylogenetic signal (right) under different scenarios. Parameters: $\sigma_{pr} = 0.25, \sigma_{hu} = 0.7, \frac{c_R}{1-\sigma_{hu}} = 0.25, (1-C) = 0.1, \beta = 3, K = 20,000, S_0 = 10,000.$

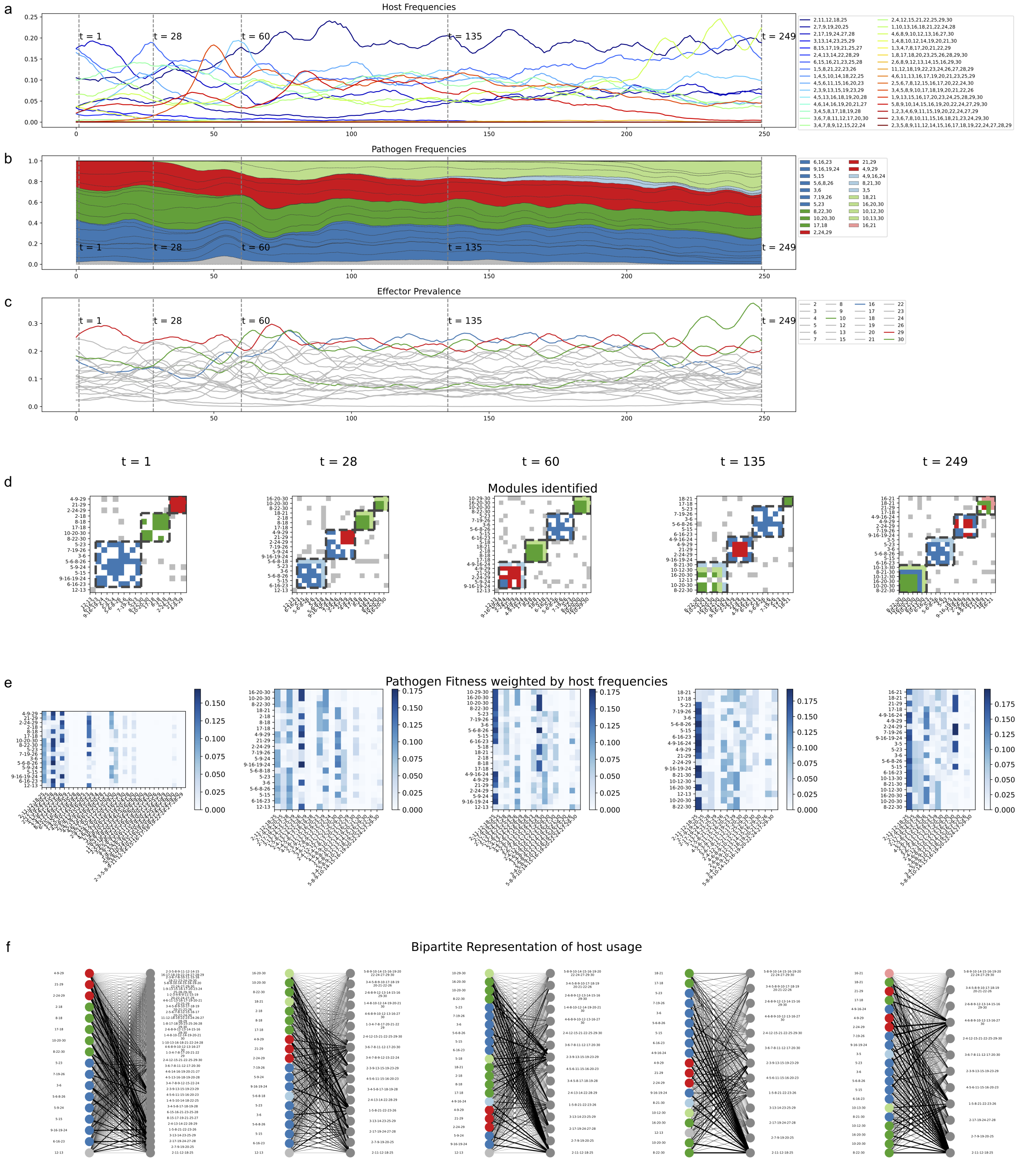


Supplementary Figure 10: The perseverance and reorganization of strain structure in the presence of introduced recombinant strains. The figure shows the first 250 generations of simulating with initial condition IC1. Otherwise similar to Supplementary FigureS 3, the strains are colored by the module membership at the initialization of the simulation, and the newly introduced strains (light blue, light green, light red) are colored based on the module membership of their parent strains (blue, green, red, respectively). All strains present at $t = 1, 28, 60, 135, 249$ are shown, regardless of their frequencies. Parameters: $\sigma_{pr} = 0.25, \sigma_{hu} = 0.7, \frac{c_R}{1-\sigma_{hu}} = 0.25, (1 - C) = 0.1, \beta = 3, K = 20,000, S_0 = 10,000.$

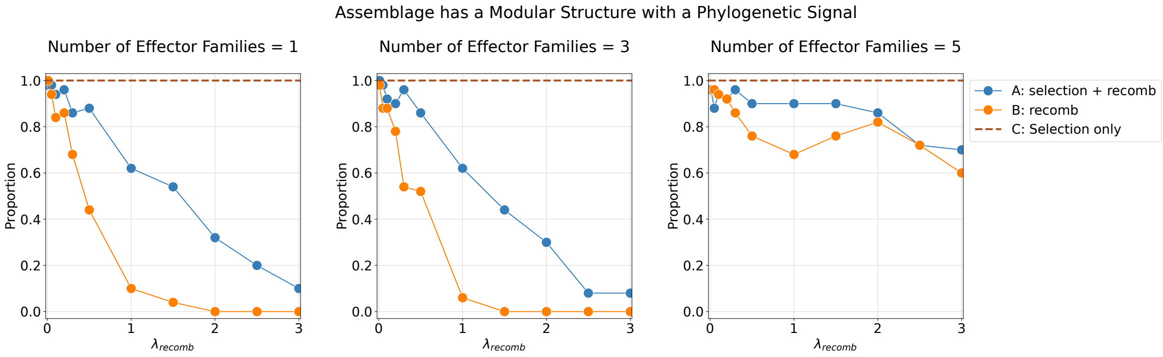


Supplementary Figure 11: The maintenance of phylogenetically aligned clusters is affected by genetic exchange restrictions due to effector similarities. Lines represent the proportion of assemblages obtained after simulating for 100 generations with a modular structure and a phylogenetic signal under different scenarios, as shown in the legend. Genetic exchange is restricted to the effectors within the defined groups with similar sequences. Results on grouping effectors into 1 (left, same as Figure 4), 3 (middle), and 5 (right) families are shown. Parameters: $\sigma_{pr} = 0.25, \sigma_{hu} = 0.7, \frac{c_R}{1-\sigma_{hu}} = 0.25, (1-C) = 0.1, \beta = 3, K = 20,000, S_0 = 10,000$.

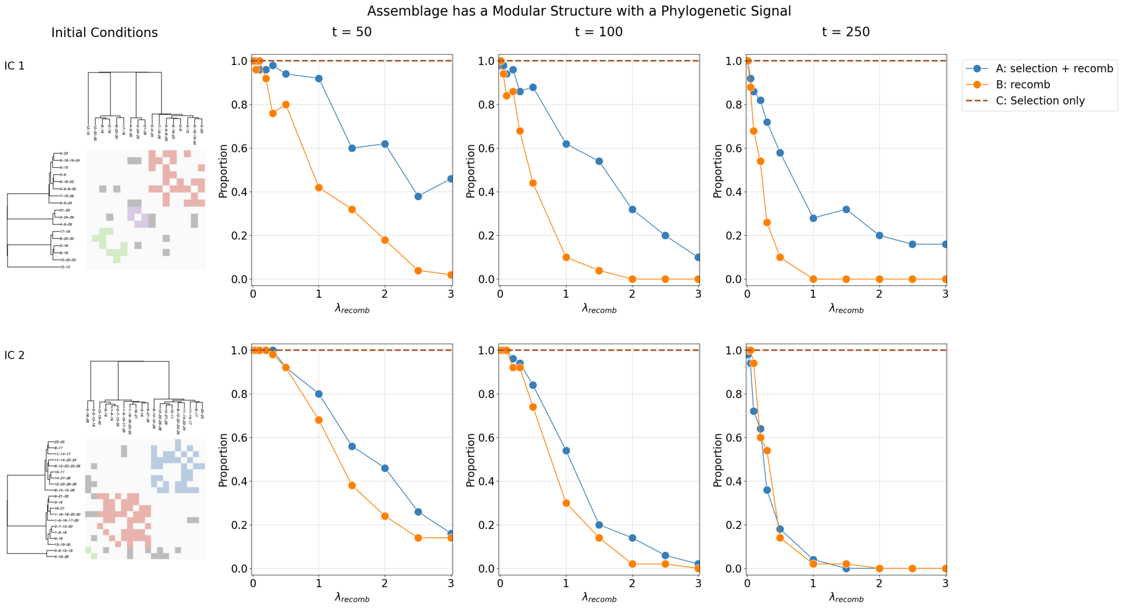


Supplementary Figure 12: The robustness of the NFDS-maintained structure through time may vary based on initial conditions. Initial conditions are shown in the left column. Lines represent the proportion of assemblages obtained at the end of the simulation with a modular structure with a phylogenetic signal at different generation numbers (50, 100, 250, as shown in the figure subtitles) under different modeling assumptions. Parameters: $\sigma_{pr} = 0.25, \sigma_{hu} = 0.7, \frac{c_R}{1-\sigma_{hu}} = 0.25, (1-C) = 0.1, \beta = 3, K = 20,000, S_0 = 10,000$.

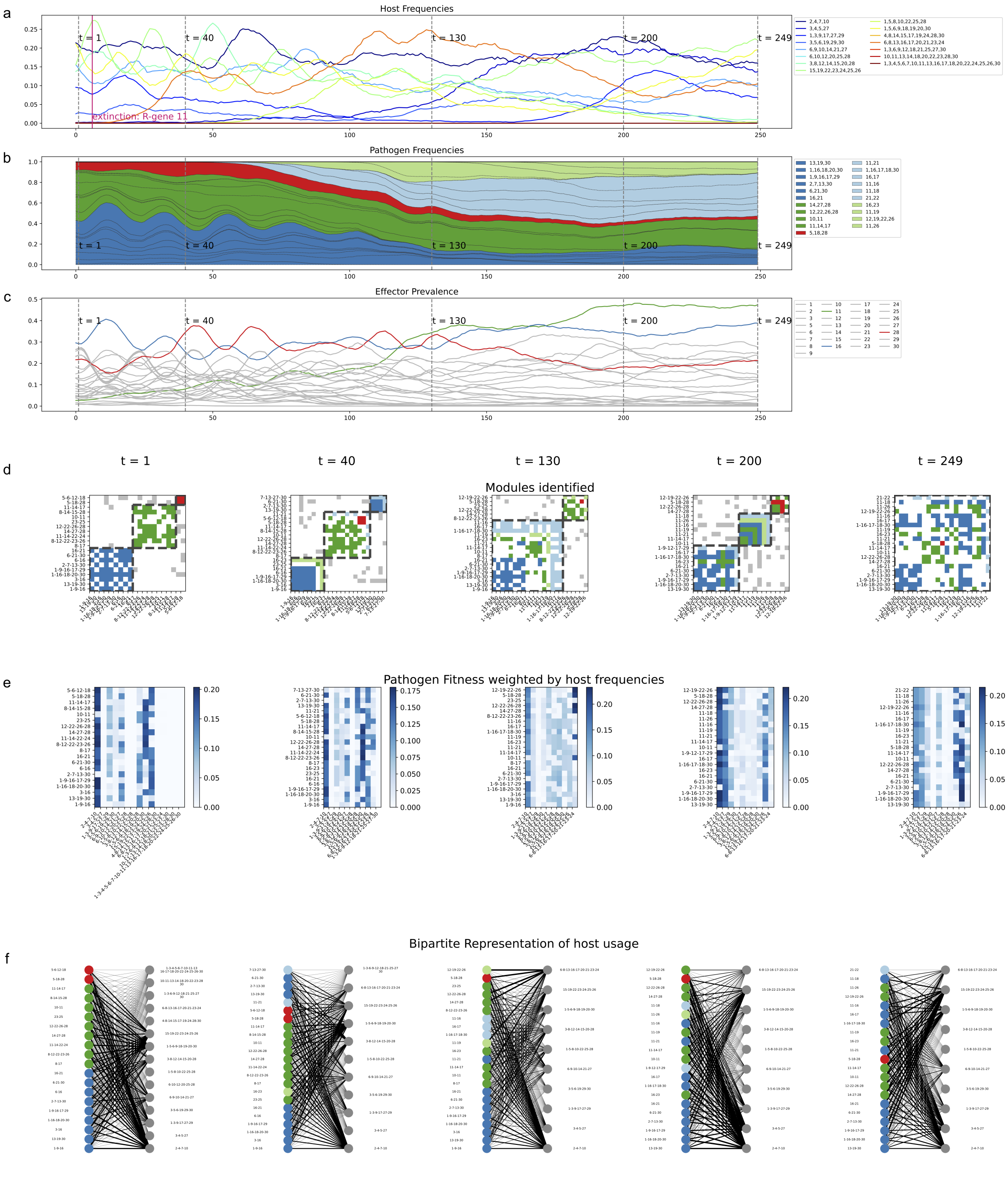


Supplementary Figure 13: The loss of strain structure over time due to recombinants. The figure shows the first 250 generations of simulating with initial condition IC2. Otherwise similar to Supplementary FigureS 10, subplot (d) at $t = 249$ shows a non-modular structure. The extinction of R-gene 11 at the early stage of the simulation is marked by red text in (a), and is followed by the increase of effector 11 prevalence, marked by the green curve in (c). Parameters: $\sigma_{pr} = 0.25, \sigma_{hu} = 0.7, \frac{c_R}{1-\sigma_{hu}} = 0.25, (1 - C) = 0.1, \beta = 3, K = 20,000, S_0 = 10,000$.

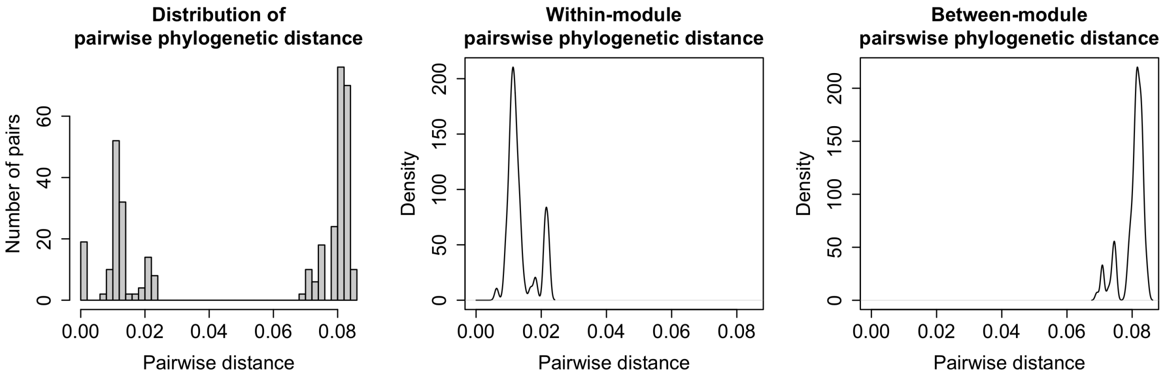


Supplementary Figure 14: The distribution of pairwise phylogenetic distances of strains from Michigan. Left: the observed distribution. Middle and Right: the estimated probability density functions for within/between module pairwise distances.

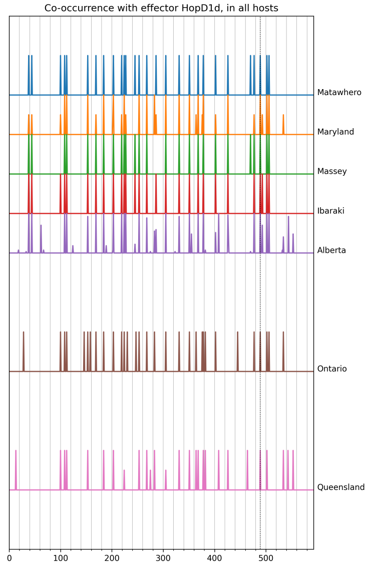


(a) HopD1d: single cluster

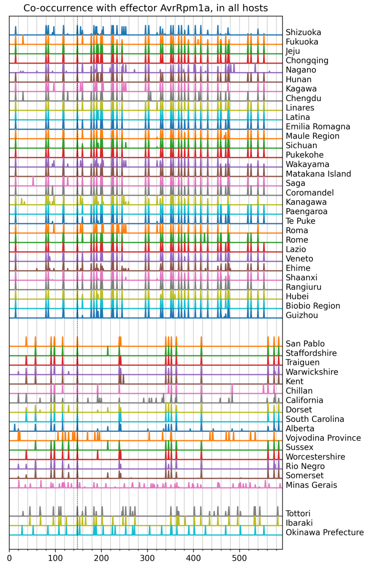


(b) AvrRpm1a: three clusters

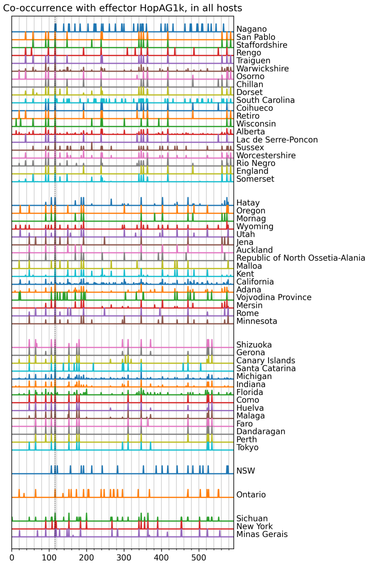


(c) HopAG1k: six clusters

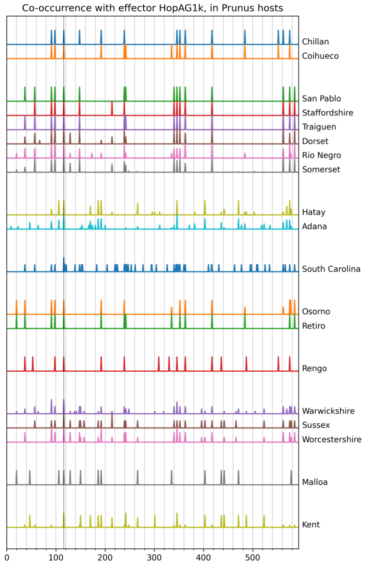


(d) HopAG1k, *Prunus* only: nine clusters

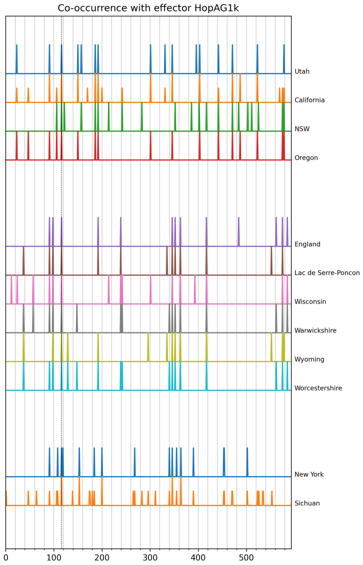


(e) HopAG1k, complete only: three clusters

Supplementary Figure 15: Examples of effector co-occurrence patterns across locations. Each plot is for a focal effector specified in the given label. The x-axis corresponds to the list of effectors and is consistent across plots (with one number per effector). Each colored line (row) represents the co-occurrence vector for the focal effector across all effectors for a given province specified in the right margin (all values have been normalized and fall in the range of 0 to 1). Identified clusters are indicated by separating the groups of rows by wider empty spaces. A co-occurrence cluster therefore reflects a group of locations with more similar patterns of co-occurrence peaks than those in other locations. For (a), although $k = 3$ clusters yield the best Silhouette score, the gap statistic suggests the clustering is not meaningful and the three clusters exhibit very similar sets of peaks. When meaningful clusters are identified, like in (b-e), there is typically no peak (or very few peaks) consistently appearing across all clusters. Thus, there are no dominant associations with particular effectors consistently found across locations. This is reflected in the existence of more than one cluster for most local effectors (see Supplementary FigureS16) (Note that the clusters in (d) and (e) are not necessarily subsets of clusters in (c). For example, in (d), the 3rd, 8th, and 9th clusters (top to bottom) belong to the second cluster in (c), while the rest belongs to the first cluster; in (e), NSW is in the same cluster as Utah, California, and Oregon, instead of forming a separate cluster as in (c). Such differences are due to changes in the density distribution in subsets of the data, which potentially lead to a stronger emphasis on more refined rather than global patterns.)

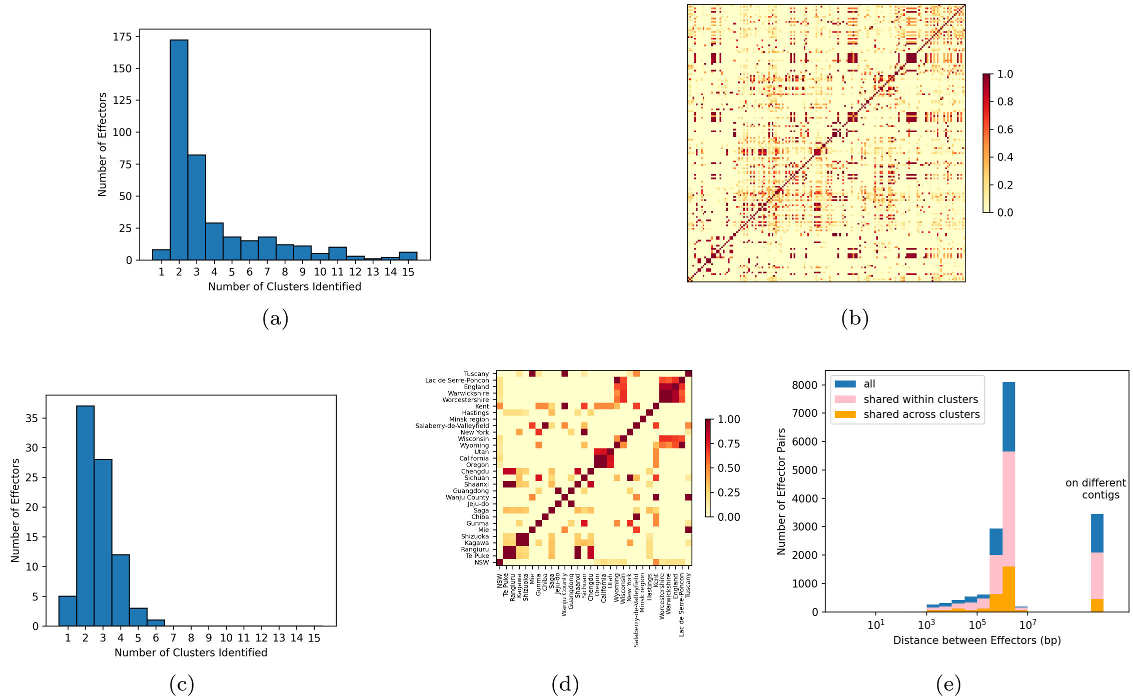


Supplementary Figure 16: A summary of co-occurrence patterns identified for all effectors across locations. (a) The distribution of the number of identified co-occurrence clusters for each focal effector. Most effectors exhibit multiple co-occurrence clusters: they are associated with different sets of effectors in different locations. (b) Proportion of effectors for which pairs of provinces fall into the same cluster, with provinces ordered by geographic proximity along the x and y axes. The low frequencies throughout the matrix illustrate that most pairs of locations do not tend to show similar patterns of co-occurrence, irrespective of geographic proximity. There are a few exceptions, with similar patterns of co-occurrence within small geographic neighborhoods (along the diagonal), and with subgroups of remote locations (off the diagonal). (The frequencies have been obtained by normalizing the number of clusters for which the two locations are found together by the total number of effectors sampled at those locations). (c) and (d) recapitulate (a) and (b) respectively but on the basis of genomes with complete assembly only (36 strains, far fewer than the 730 original ones). This restriction results in a reduced number of identified clusters, although the main features remain. (e) The distribution of pairwise effector distances for genomes with complete assembly within clusters, across clusters, and for all. Different colors differentiate these three subgroups: blue represents for each focal effector, its distance to all effectors; pink, its distance to effectors that it always co-occurs with when classified into the same cluster; orange, its distance to effectors it always co-occurs with, regardless of cluster assignations. There are no notable differences among the distributions, including the typical distances for these categories. Thus, co-occurrence patterns do not reflect physical proximity in the genome.

#### 7.1 Recombination rate estimate

| Paramter | Definition | Estimate | Source |
|---|---|---|---|
| $\frac{r}{m}$ | effective recombination rate | 8.1 | [97] |
| $\theta$ | per site mutation rate | $1.4 \times 10^{-1}$ | [97] |
| $\frac{1}{\delta}$ | inverse importation tract length | $7.8 \times 10^{-3}$ | [49] |
| $\nu$ | estimated sequence divergence | $4 \times 10^{-2}$ | [49] |
| $g$ | number of pathogen generations per year | $[1.5 \times 10^3, 3 \times 10^3]$ | [81] |
| $N_e$ | effective population size | $[10^8, 10^9]$ | [20] |
| $n_{eff}$ | number of effectors per strain | [10, 20] | see data repository 4 |
| $L_{eff}$ | average length per effector | $10^3$ | see daa repository 4 |
| $n_P$ | number of lineages | [10, 20] | Simulation setup |

Supplementary Table 1: Parameters and their estimated values for estimating the expected number of recombination events per plant generation.

We model recombination events with a Poisson process [19],

$$P(K = k|t) \sim Pois(\rho t),$$

such that the expected number of recombination events ($E[K|t]$) per site per lineage per coalescence time unit is $\rho$.

Rearranging the derivation of effective recombination rate $\frac{r}{m} = \frac{\rho\nu\delta}{\theta}$ [97], $\rho$ can be expressed as

$$\rho = \frac{\theta r}{\nu\delta m}.$$

To obtain a per host generation (per year) recombination rate, we define $g$, the number of pathogen generations per year, and therefore with a pathogen generation time of $\frac{1}{g}$ years, we rescale one year to $\frac{1}{N_e \frac{1}{g}} = \frac{g}{N_e}$ coalescence time units.

We estimate the total number of sites of interest undergoing recombination per strain with $n_{eff} \times L_{eff}$, the product of approximations of the number of effectors per strain and effector lengths.

Finally, we multiply the per lineage per year recombination rate with $n_P$, the number of lineages, to obtain the total recombination rate of all simulated strains. Therefore, the total expected number of recombination can be expressed as

$$
\begin{aligned}
n_{eff} \cdot L_{eff} \cdot n_P \cdot E[K|t = 1yr] &= n_{eff} \cdot L_{eff} \cdot n_P \cdot \rho \cdot \frac{g}{N_e} \\
&= n_{eff} \cdot L_{eff} \cdot n_P \cdot \frac{\theta r}{\nu \delta m} \cdot \frac{n}{N_e} \\
&= n_{eff} \cdot L_{eff} \cdot n_P \cdot \frac{r}{m} \cdot \theta \cdot \frac{1}{\delta} \cdot \frac{1}{\nu} \cdot \frac{n}{N_e},
\end{aligned}
$$

yielding an estimate with a lower bound of $3.3 * 10^{-2}$ and an upper bound of 2.7 (recombination events per year). The parameters and estimates are summarized in Table S 1.
